# Where the Minor Things Are: A Pan-Eukaryotic Survey Suggests Neutral Processes May Dominate Minor Spliceosomal Intron Evolution

**DOI:** 10.1101/2022.09.24.509304

**Authors:** Graham E. Larue, Scott W. Roy

## Abstract

Spliceosomal introns are gene segments removed (“spliced”) from RNA transcripts by large ribonucleoprotein machineries called spliceosomes. In some eukaryotes a second spliceosome (the minor/ U12-type) is responsible for processing a tiny minority of introns. Despite its seemingly modest role, minor splicing has persisted for roughly 1.5 billion years of eukaryotic evolution. Identifying and cataloging minor introns in > 3000 eukaryotic genomes, we report diverse evolutionary histories including surprisingly high numbers of minor introns in some fungi and green algae, repeated massive loss, as well as several general biases in the positional and genic distributions of minor introns. We estimate that ancestral minor intron densities were comparable to those of the most minor intron-rich species, suggesting a trend of long-term stasis. Finally, three findings suggest a major role for neutral processes in minor intron evolution. First, we find highly similar patterns of minor and major intron evolution, in contrast to the predictions of both functionalist and deleterious models. Second, we find that observed functional biases among minor intron-containing genes are largely explained by these genes’ greater ages. Third, we find no association of intron splicing with cell proliferation in a minor intron-rich fungus, suggesting that regulatory roles are lineage-specific and thus cannot offer a general explanation for minor splicing’s persistence. These data constitute the most comprehensive view to date of modern minor introns, their evolutionary history, and the forces shaping minor splicing, and provide a foundation for future studies of these remarkable genomic elements.

## 1 Introduction

Spliceosomal introns are sequences in eukaryotic genes that are removed (spliced) from the pre-mRNA transcripts of genes by machinery known as the spliceosome prior to maturation and nuclear export of the final mRNA [92, 51, 37]. For the better part of a decade after spliceosomal introns (hereafter simply introns) were first characterized in eukaryotic genomes [36, 29, 7, 8, 60], it was assumed that all introns shared a fixed set of consensus dinucleotide termini—GT at the beginning (5^*′*^ side) and AG at the end (3^*′*^ side)—and were processed by the same core machinery [54, 34]. This view was revised after the discovery of a small number of introns with AT-AC termini [34, 30], and shortly thereafter an entirely separate spliceosome was described that could process these aberrant introns [89, 88], termed the U12 or minor spliceosome. The minor spliceosome, like its counterpart now known as the major/U2 spliceosome, has origins early in eukaryotic evolution [73]. Since minor introns were first documented as having AT-AC termini, it has been shown that the majority of minor introns in most species are in fact of the GT-AG subtype, although an increasing diversity of termini (so-called “non-canonical” introns, with boundaries that aren’t GT-AG, GC-AG or AT-AC) seem to be able to be processed in certain contexts and to varying degrees by both spliceosomes [25, 12, 64, 81, 56]. Until very recently [41], in every genome investigated minor introns have been found to comprise only a tiny fraction (<∼0.05%) of the total set of introns; despite this, they have also been found to be well-conserved over hundreds of millions of years of evolution (e.g., 96% of minor introns in human are conserved in chicken, 83% in octopus (Fig. 2)).

While minor introns were almost certainly present in early eukaryotes [73] and are retained in a wide variety of eukaryotic lineages [48, 56], to date only two lineages—animals and plants —are known to retain more than a few dozen minor introns[56]. Interestingly, in contrast to the massive minor intron loss observed in many lineages, minor intron complements in certain clades are remarkably evolutionarily stable. This contrasting pattern of retention versus massive loss raises a puzzle of minor spliceosomal intron function: if minor introns are not functionally important why are they almost entirely conserved over hundreds of millions of years in some lineages; yet if they are important, how can they be repeatedly decimated or lost entirely in other lineages?

Two observations are particularly relevant to the question of minor spliceosomal intron function. First, over the past ten or so years, some studies have proposed roles for minor introns in cellular differentiation, with decreased minor splicing activity driving downregulation of minor intron-containing genes (MIGs) associated with cessation of cell cycling [27, 22, 39]. Most compellingly, a recent study showed that the splicing regulator SR10 is regulated at the level of minor splicing, with inefficient splicing leading to downregulation of other SR proteins whose pro-splicing activities promote cell cycle progression [52]. Interestingly, a negative association of minor splicing with cell differentiation has also been argued for in plants [27, 2]. This pattern is curious, given that the common ancestor of animals and plants is thought to have been unicellular and thus not to have undergone terminal differentiation (although recent findings of multicellular stages as well as differentiation-like processes across diverse eukaryotes may ultimately call this common assumption into question [57, 9]).

Second, minor introns have been reported to show functional biases, being disproportionately found in genes encoding various core cellular functions including DNA repair, RNA processing, transcription and cell cycle functions that largely appear to hold between plants and animals [27, 11] (though the strength of these associations is questioned somewhat in [5]). Particularly given the above evidence that regulation of minor splicing regulates core cellular processes, these patterns would seem to be consistent with an ancient role for minor splicing in cell cycle regulation, that could have been secondarily recruited for multicellular differentiation separately in animals and plants. However, other explanations remain possible. What is needed to understand the evolutionary history and importance of minor spliceosomal introns is genomic and regulatory characterization of additional lineages with relatively large complements of minor spliceosomal introns.

Because minor introns possess sequence motifs distinct from major introns [10, 11], it is possible to identify them using sequence-based bioinformatic methods [56, 1, 11, 3, 80, 43]. Previous studies have cataloged the presence/absence of minor introns/spliceosome components across multiple eukaryotic genomes using various custom tools, [48, 3, 80, 1], but many of these studies were necessarily constrained in their analyses by the limits of the data available at the time, and by the lack of a published or otherwise convenient computational method to identify minor introns.

In this work, with the substantially larger and more diverse genomic datasets now publicly accessible coupled with the intron classification program `intronIC` [56], we have been able to aggregate minor intron presence/absence data with higher fidelity than earlier works across a much larger sample of eukaryotic species. By compiling a catalogue of minor introns across thousands of eukaryotic species, we provide an unprecedentedly general portrait of minor intron characteristics, diversity, and distribution, and test several important hypotheses about minor intron evolution and function. In addition, we use the discovery of hundreds of minor spliceosomal introns in a mycorrhizal fungus to characterize several features of the minor spliceosomal system across cell types, and suggest that the functional biases long observed in minor introns may largely be explained by the age bias of their parent genes.

## 2 Methods

### 2.1 Data acquisition

Genomes and annotations for 3,108 eukaryotic species were downloaded from the online databases hosted by NCBI (RefSeq and GenBank), JGI and Ensembl using custom Python scripts, and taxonomic information for each species was retrieved from the NCBI Taxonomy Database [24] using a Python script. We used NCBI as our primary resource, since it contains the largest number of species and in many cases serves as the upstream source for a number of other genome resources. Additional species available only from JGI and Ensembl were included for completeness, as were GenBank genomes with standard annotation files (GTF or GFF; species with only GBFF annotations were excluded). GenBank annotations in particular are of highly variable quality and may be preliminary, draft or otherwise incomplete, which is one of the reasons we chose to also include annotation `BUSCO` scores for all species. Accession numbers (where available) and retrieval dates of the data for each species are available under the following DOI: https://doi.org/10.6084/m9.figshare.20483655.

### 2.2 Identification of spliceosomal snRNAs

Each genome was searched for the presence of the minor snRNAs U11, U12, U4atac and U6atac using `INFERNAL` v1.3.3 [58] with covariance models retrieved from Rfam (RF00548, RF00007, RF00618, RF00619). The default E-value cutoff of 0.01 was used to identify positive snRNA matches, and any snRNA with at least one match below the E-value cutoff was considered present.

### 2.3 Classification of minor introns

For every genome with annotated introns, `intronIC` v1.2.3 [56] was used to identify putative minor introns using default settings which includes introns defined by exon features (e.g., introns in UTRs) and excludes any introns shorter than 30 nt. Although our substrate data includes UTR introns — at least some of which appear to be involved in the regulation of gene expression [6, 16, 84, 79, 50] — the analyses performed in this study include only introns in protein-coding regions of genes. UTR introns generally are an understudied class of introns, and almost nothing is known about minor introns in UTRs; exploring this in more detail would certainly be an exciting avenue for future research.

### 2.4 Identification of orthologous introns

A set of custom software was used to identify orthologs between various species as described previously [41]. Briefly, transcriptomes for each species in a group were generated using the longest isoforms of each gene (https://github.com/glarue/cdseq). Then, the program `reciprologs` (https://github.com/glarue/reciprologs) was used in conjunction with `DIAMOND` v2.0.13 (flags: `--very-sensitive --evalue 1e-10`) to identify reciprocal best hits (RBHs) between all pairs of species, and to construct an undirected graph using the RBHs as edges. The maximal cliques of such a graph represent orthologous clusters where all members are RBHs of one another. In certain cases, when only orthologous MIGs were required (as opposed to all orthologs), `reciprologs` was run as part of a pipeline using the `--subset` argument in combination with separately generated lists of MIGs for each species, which dramatically decreases runtime by constraining the query lists to only include MIGs (producing identical results to the subset of a full alignment containing MIGs). Full ortholog searches were required for all ancestral density reconstructions as well as all comparisons of minor and major intron conservation/loss (e.g., Fig. 5a).

Orthologous groups were aligned at the protein level using a combination of `MAFFT` v7.453 and `Clustal Omega` v1.2.4, and intron positions within the alignments were computed using a custom Python pipeline (following the approach in [68]). Local alignment quality of ≥ 40% conserved amino acid identity (without gaps) over a window of 10 residues both upstream and downstream of each intron position was required. Introns of the same type in the same position within orthologs so aligned were considered conserved. For the analyses of putative intron type conversions (e.g., minor-to-major), major introns were required to have scores ≤ −30 instead of the default threshold of 0 to minimize the inclusion of minor introns with borderline scores as major-type, and intron alignments containing introns with such borderline scores (a tiny fraction of the total alignments) were excluded. Intron sliding (the shifting of an individual intron’s boundaries within a gene versus its ancestral location) [85] is not explicitly accounted for by our pipeline (an intron sliding event would be categorized as intron loss in the containing gene); however, this phenomenon is at most very rare and unlikely to meaningfully affect our results [63, 67, 85, 77].

### 2.5 Intron position within transcripts and intron phase

Included in the output of `intronIC` [56] is information about the relative position of each intron within its parent transcript, represented as a percentage of the total length of coding sequence, as well as intron phase (for introns defined by CDS features). These were extracted for every species and used in the associated analyses.

### 2.6 Non-canonical minor introns

Species were first analyzed to assess the number of putative non-canonical minor introns they contained, and those with the highest numbers of non-canonical minor introns were used to perform multiple alignments of different pairs of species. From these alignments, all orthologous intron pairs with a minor intron (minor-minor or minor-major) were collected, and used to build clusters (subgraphs) of orthologous introns. For animals, human was used in a majority of the alignments to facilitate the generation of larger subgraphs (where the same intron shared between different pairwise alignments will group the alignments together); for plants, *Elaeis guineensis* served the same purpose.

### 2.7 BUSCO analysis

Translated versions of the transcriptomes of all species were searched for broadly-conserved eukaryotic genes using `BUSCO` v5.3.2 [82] and the `BUSCO` lineage `eukaryota_o db10`. Complete `BUSCO` scores were then integrated into the overall dataset (e.g., Fig. 1 and Fig. 4).

**Fig. 1.**
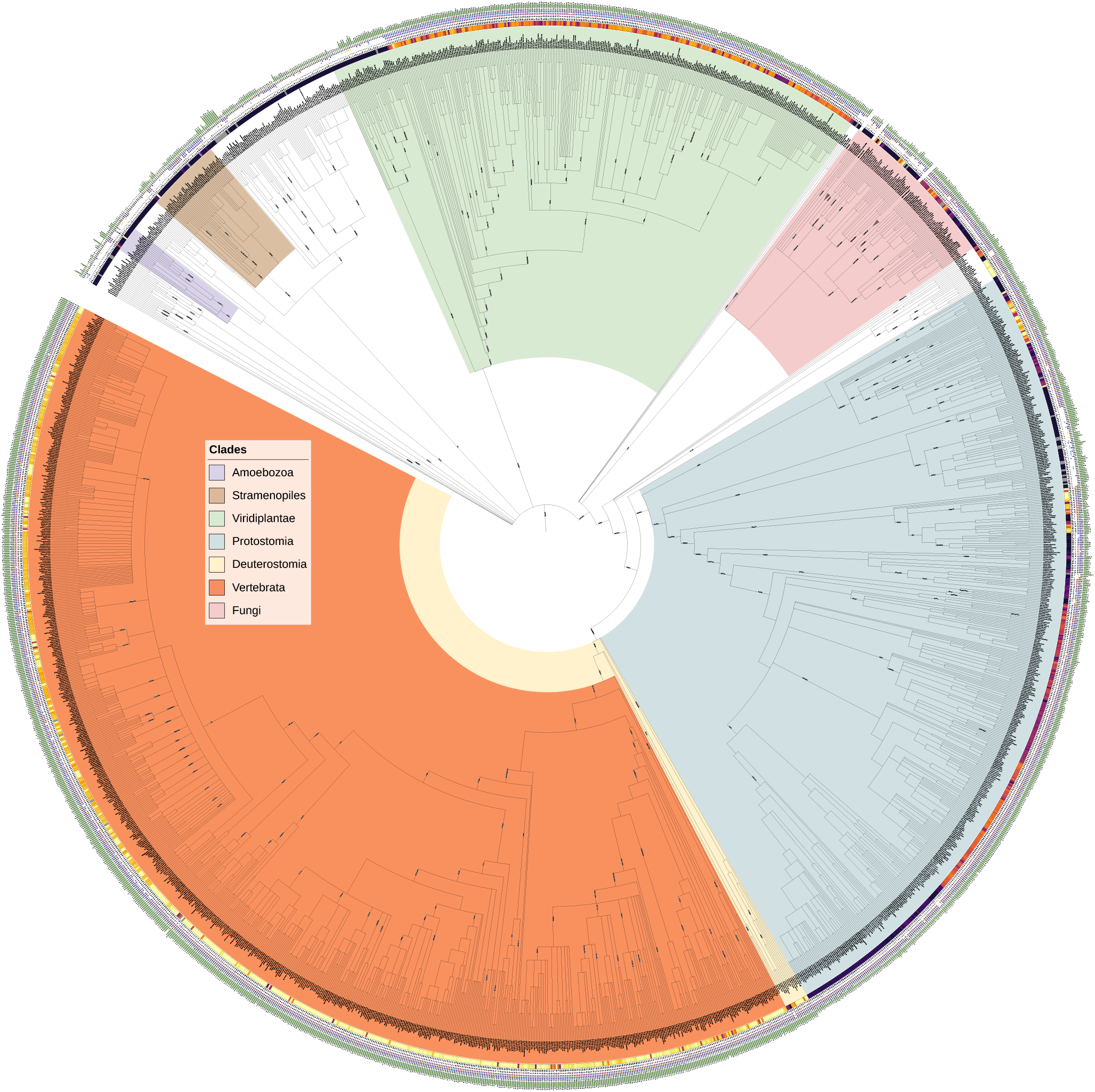
Minor intron densities for thousands of eukaryotic species. The colored strip following the species name represents the relative minor intron density (darker = lower, brighter = higher, gray values indicate species for which the estimated values are less confident, and may be enriched for false positives; see Methods). Additional data from inside to outside is as follows: minor intron density (%), number of minor introns, presence/absence of minor snRNAs in the annotated transcriptome (red: U11, light blue: U12, yellow: U4atac, purple: U6atac), `BUSCO` score versus the eukaryotic `BUSCO` gene set, average overall intron density in introns/kbp coding sequence. Taxonomic relationships based upon data from the NCBI Taxonomy Database [24]; figure genera.ted using iTOL [42]

### 2.8 Curation of minor intron data/edge cases

Due to the number of genomes involved in our analyses, there may be some number of introns that appear superficially similar to minor introns simply by chance, and `intronIC` is unable to filter out such introns because it does not account for additional factors like local context, presence/absence of snRNAs, etc. In general this is not an issue, as the number of false-positive minor introns per genome is usually very small. However, when summarizing aggregate eukaryotic data and attempting to provide a resource to be used as a reference, we felt that some amount of curation was warranted to avoid the inclusion of obviously spurious results.

We therefore used the following heuristics in deciding whether to designate a given species as either confidently containing or not containing minor introns—species not meeting either set of criteria were assigned a minor intron density color of gray in Fig. 1. First, it is important to note that `intronIC` will automatically try to adjust intron boundaries by a short distance if the starting boundaries are non-canonical and there is a strong minor 5^*′*^SS motif within ∼10 bp of the annotated 5^*′*^SS. In some poorly-annotated species, or species with otherwise aberrant intron motifs this can lead to increased false positives in the form of putatively minor introns with “corrected” splice boundaries. Such introns are documented by `intronIC` in the output, so it is possible to determine their proportion in the final number of minor introns reported. The criteria for presence of minor introns in our dataset is a corrected minor intron fraction of ≤ 0.25, ≥ 3 called minor introns and at least two minor snRNAs.

The criteria for absence of minor introns (assigned the color black in the minor intron density color strip in Fig. 1) is either of the following: ≤ 3 called minor introns and fewer than two minor snRNAs; ≤ 5 called minor introns and fewer than two minor snRNAs and fewer than five uncorrected ATAC minor introns and either annotated in RefSeq or with a `BUSCO` score greater than or equal to *B*_*Q*1_− (1.5 *× B*_*IQR*_), where *B* is all the `BUSCO` scores of the broad RefSeq category to which the species belongs (i.e., “vertebrates”, “invertebrates”, “plants”, “protozoa”, “fungi”), *B*_*Q*1_ is the first quartile of such scores and *B*_*IQR*_ is the inner quartile range of such scores. The idea behind this metric is to only assign confident minor intron loss to species whose `BUSCO` scores aren’t extremely low; very low `BUSCO` scores could indicate real gene loss or incomplete annotations, and neither of those scenarios forecloses on the possibility that the species may have minor introns (whereas a species with a high `BUSCO` score and a very low number of minor introns/minor snRNAs is more likely to be genuinely lacking either/both). Finally, species with very low numbers of minor introns and minor snRNAs but very high minor intron densities (≥ 1%) were categorized as uncertain to account for a small number of edge cases with massive intron loss and spurious false positives that, due to the low number of total introns, misleadingly appear to be cases of outstandingly high minor intron density (e.g., *Leishmania martiniquensis*). Importantly, Fig. 1 still includes the raw values for each species matching the above criteria; it is only the minor intron density color which is adjusted to indicate lack of confidence.

### 2.9 Calculation of summary statistics (introns/kbp CDS, transcript length, etc.)

Transcriptomes for all species were generated using a custom Python script (https://github.com/glarue/cdseq). Briefly, each annotated transcript’s length was calculated as the sum of its constituent CDS features, and the longest isoform for each gene was selected. The number of introns per transcript was computed based on the same data, and combined with the transcript length to calculate introns/kbp coding sequence for each gene. Intron lengths were extracted directly from `intronIC` output, as was intron phase and intron position as a fraction of transcript length (where the position of each intron, taken as the point position in the coding sequence where the intron occurs, is calculated as the cumulative sum of the preceding coding sequence divided by the total length of coding sequence in the transcript). For comparisons of intron densities and gene lengths of MIGs and non-MIGs, species with fewer than ten putative minor introns were excluded to avoid inclusion of spurious minor intron calls.

### 2.10 Ancestral intron density reconstruction

Reconstructions of ancestral intron complements in different nodes was performed as described in [69]. Briefly, for a set of three species *α, β* and *γ* where *γ* is an outgroup to *α* and *β* (i.e., *α* and *β* are sister with respect to *γ*), introns shared between any pair of species are (under the assumption of negligible parallel intron gain) *a priori* part of the set of introns in the ancestor of *α* and *β*. For all introns shared between a given species pair, for example *α* and *γ* (but not necessarily *β*) *N*_*αγ*_, the probability of an intron from that set being found in *β* (in other words, the fraction of ancestral introns retained in *β*) is

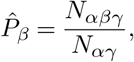

where *N*_*αβγ*_ is the number of introns shared between all three species. Deriving these fractions of ancestral introns for each of the aligned species, we then define *N*_Ω_ as the total number of ancestral introns, and its relationship to the conservation states of introns in the alignments of the three species as

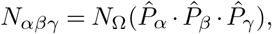

the product of the ancestral intron number and the fraction of ancestral introns present in each species. Finally, solving for the number of ancestral introns we get the estimate

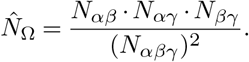

Performing the above procedure for both major and minor introns in a given alignment allowed us to estimate the ancestral minor intron density for the corresponding node as

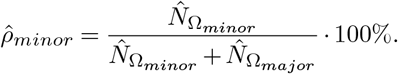

However, without some point of reference 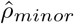 is difficult to interpret, as the genes included in the alignments are not an especially well-defined set—because these genes are simply all of the orthologs found between a given trio of species, their composition is likely to change at least somewhat for each unique group of aligned species representing the same ancestral node. We dealt with this by normalizing to a chosen reference species included in each group. For example, in our reconstructions of intron densities in the ancestor of Diptera, human was used as the outgroup and was therefore present in all alignments. After calculating the estimated minor intron density in the Dipteran ancestor, we then divided that value by the minor intron density in the human genes present in the same alignments to produce the estimated ancestral minor intron density relative to the corresponding minor intron density in human. Because using human as the outgroup for reconstructions of fungal and plant ancestors results in very small absolute numbers of minor introns, kingdom-specific outgroups were chosen instead: the estimates of ancestral fungal densities are relative to *Rhizophagus irregularis*, and those for plants are relative to *Lupinus angustifolius*. Because multiple species combinations were used to estimate the minor intron density at each ancestral node, we report the mean value over all *n* estimates for each node

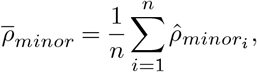

± the standard error (Fig. 10).

As has been pointed out in other contexts, ancestral state reconstructions may be confounded by several different factors [32, 23, 18]. Many such concerns are minimized in our specific application given that a) the traits under consideration are not complex, but rather the simple binary presence/absence of discrete genetic elements, b) calculations are restricted to introns present in well-aligning regions of orthologs (thereby avoiding issues with missing gene annotations in a given species, since alignments must include sequences from all species to be considered) and c) the contribution of parallel intron gain, especially of minor introns, is likely to be very small [71, 86, 15]. There are a number of other potential sources of bias in our analyses, however, which are worth addressing. First, our ancestral intron density estimates are (to a large, though not complete, extent) dependent upon the accuracy of the phylogenetic relationships in Fig. 1. Ideally, we would have perfect confidence in all of the relationships underlying each node’s reconstruction, but such an undertaking is beyond both the scope of this paper and the expertise of its authors. While we have done our best to be assiduous in choosing nodes with well-resolved local phylogenies—which is one reason we have not provided similar reconstructions for a much larger number of nodes with less-confident phylogenetic relationships—it remains the case that our reconstructions are only fully informative with respect to the tree upon which they are based. That being said, unless the phylogeny for a given node is so incorrect as to have mistaken one of the ingroups for the outgroup (i.e., the chosen outgroup was not in fact an outgroup), the reconstruction should still represent the ancestor of the two ingroup species. Second, we are relying on the correct identification of minor introns within each species to allow us to identify conserved/non-conserved minor introns in multi-species alignments. Although the field in general lacks a gold-standard set of verified minor introns upon which to evaluate classifier performance, the low empirical false-positive rate of `intronIC` (as determined by the number of minor introns found in species with compelling evidence for a lack of minor splicing) and the high degree of correspondence of its classifications with previously-published data suggests that our analyses are capturing the majority of the minor introns in each alignment. There is also the possibility that many minor introns are unannotated in many genomes (and in fact, for certain annotation pipelines we know that this has historically been the case). This concern is mediated somewhat by the fact that, because we are only considering gene models that produce well-aligning protein sequences across multiple species, our alignments are unlikely to contain unannotated introns of either type. Unannotated minor introns, necessarily residing in completely unannotated genes, would of course not be considered in our analyses, which would result in a shrinking of the total number of orthologous genes compared. Due to the law of small numbers, this could raise concerns that the chosen samples may not reliably represent the complete data with sufficient confidence. We have done what we can to combat this by choosing species with annotations of high quality (as assessed by `BUSCO` completeness, for example), and by using multiple combinations of species to reconstruct each node—for reconstructions based upon a large number of different alignments, the low standard errors of the estimates give us some confidence that this kind of missing data is unlikely to qualitatively change our results.

### 2.11 Differential gene expression

Single-end RNA-seq reads from previously-published celltype-specific sequencing of *Rhizophagus irregularis* [38] (four biological replicates per cell type) were pseudoaligned to a decoy-aware version of the transcriptome using `Salmon` v1.6.0 [62] (with non-default arguments `--seqBias --s oftclip`). The `Salmon` output was then formatted with `tximport` v1.14.2 [83], and differential gene expression (DGE) analysis was performed using `DESeq2` v1.26.0 [49] with the following arguments: `test=“LRT”`, `useT=TRU E`, `minReplicatesForReplace=Inf`, `minmu=1e-6`, `reduced=` ∼`1`. For each pairwise combination of cell types, genes with significant DGE values (Wald p-value < 0.05) were retained for further analysis.

### 2.12 *Rhizophgaus* z-score metric

Following the methodology used by Sandberg et al. to assign a proliferation index to cell types [76], z-scores were calculated per feature (whether for gene expression or intron retention/splicing efficiency) across all cell-type replicates (n=20), and then summarized for each cell type by the mean value of the corresponding replicate z-scores (departing from the reference method in this aspect). Prior to conversion to z-scores, the raw gene expression data was normalized by running the output from `tximport` through the `fpkm()` function in `DESeq2`. For group z-score comparisons (e.g., proliferation-index genes, minor introns vs. major introns), the median of the top 50% of z-scores from each group was used. As the z-score calculation requires there to be variation across samples, certain genes/introns were necessarily omitted under this metric.

### 2.13 Intron retention and splicing efficiency

For each RNA-seq sample, `IRFinder-S v2.0` [53] was used to compute intron retention levels for all annotated introns. Introns with warnings of “LowSplicing” and “Low-Cover” were excluded from downstream analyses. Across replicates within each cell type, a weighted mean retention value was calculated for each intron, with weights derived by combining the average number of reads supporting the two intron-exon junctions and the total number of reads supporting the exon-exon junction.

Intron splicing efficiency was calculated as previously described [41]. Briefly, RNA-seq reads were mapped to splice-junction sequence constructs using `Bowtie` v1.2.3 [40] (excluding multiply-mapping reads using the non-default argument `-m 1`). Introns with fewer than five reads supporting either the corresponding exon-exon junction or one of the intron-exon junctions (or both) were excluded. For each intron, the proportion of reads mapped to the intron-exon junction(s) versus the exon-exon junction was used to assign a splicing efficiency value for each sample (see reference for details). Within each cell type, the weighted mean of replicate splicing efficiency values for each intron was calculated in the same manner as for intron retention.

### 2.14 Spliceosome-associated gene expression

Orthologs of human spliceosome components were found in *Rhizophagus irregularis* via a reciprocal-best-hit approach (https://github.com/glarue/reciprologs) using `BLAST` v2.9.0+ [14] with an E-value cutoff of 1 *×* 10^−10^. Four genes from each splicing system (major and minor) were identified in *Rhizophagus* by this approach, consisting of orthologs to human minor spliceosome genes ZMAT5 (U11/U12-20K), RNPC3 (U11/U12-65K), SNRNP35 (U11/U12-35K), and SNRNP25 (U11/U12-25K) and major spliceosome genes SF3A1 (SF3a120), SF3A3 (SF3a60), SNRNP70 (U1-70K) and SNRPA1 (U2 A^*′*^). Gene expression values generated by `Salmon` for each set of genes in each cell type were averaged across replicates, and pairwise comparisons between cell types were made for the same set of genes (e.g., minor spliceosome genes in IS vs. MS). The significance of differences in expression between paired gene sets from different cell types was assessed using a Wilcoxon signed-rank test, with p-values corrected for multiple testing by the Benjamini-Hochberg method.

## 3 Results

### 3.1 Minor intron diversity in thousands of eukaryotic genomes

In order to better assess the landscape of minor intron diversity in eukaryotes, we used the intron classification program `intronIC` [56] to process ∼270 million intron sequences and uncover minor intron numbers for over 3000 publicly-available eukaryotic genomes, representing to our knowledge the largest and most diverse collection of minor intron data assembled to date (Fig. 1, underlying plain text data available at https://doi.org/10.6084/m9.figshare.20483655).

Of the 1844 genera represented in our data, 1172 (64%) have well-supported evidence of minor introns in at least one species (see Methods for details), while the remaining 672 appear to lack minor introns in all available constituent species. Consistent with previous studies [11, 3, 35, 80, 1, 87, 44, 56, 90], minor intron numbers and densities (fractions of introns in a given genome classified as minor type) vary dramatically across the eukaryotic tree; average values are highest in vertebrates and other animals, while variation between species appears to be lowest within land plants. Conservation of minor introns between different pairs of species is largely consistent with previously-published results [1, 3, 44, 56] (Fig. 2). The intriguing pattern of punctuated wholesale loss of minor introns is apparent within many larger clades in our data, along with a number of striking cases of minor intron enrichment in otherwise depauperate groups.

**Fig. 2.**
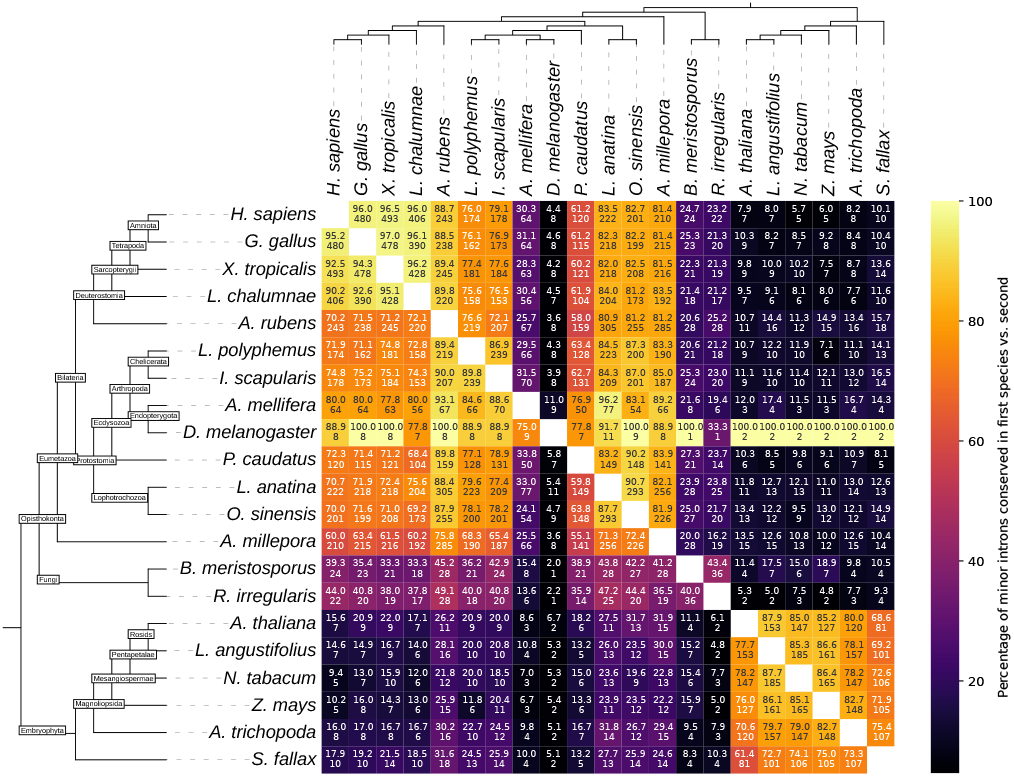
Pairwise minor intron conservation between various species. Bottom number is the number of minor introns conserved between the pair; top number is the number of conserved minor introns as a percentage of the minor introns present in the alignments for the associated species (the row species). For example, there are eight minor introns conserved between *D. melanogaster* and *L. polyphemus*, which is 88.9% of the *Drosophila* minor introns present in the alignment, but only 4.3% of *Limulus* minor introns. Full names of species are as follows: *Homo sapiens, Gallus gallus, Xenopus tropicalis, Latimeria chalumnae, Asterias rubens, Limulus polyphemus, Ixodes scapularis, Apis mellifera, Drosophila melanogaster, Priapulus caudatus, Lingula anatina, Octopus sinensis, Acropora millepora, Basidiobolus meristosporus, Rhizophagus irregularis, Arabidopsis thaliana, Lupinus angustifolius, Nicotiana tabacum, Zea mays, Amborella trichopoda, Sphagnum fallax*

#### 3.1.1 Minor intron enrichment

A number of cases of minor-intron-rich lineages are worth highlighting. As shown in Fig. 3, the highest known minor intron density is found within the Amoebozoa; our recently-reported data in the slime mold *Physarum polycephalum* [41] dwarfs all other known instances of local minor intron enrichment and appears to be an extremely rare example of significant minor intron gain. In the present study, we also find relatively high numbers of minor introns (compared to other amoebozoan species) in both the flagellar amoeba *Pelomyxa schiedti* (n=90) and the variosean amoeba *Protostelium aurantium* (labeled *Planoprotostelium fungivorum*^1^ in Fig. 1) (n=265). Although the numbers of minor introns in these species conserved with minor introns in other lineages (e.g., human) are very low, in all cases we find at least some degree of conservation. For example, in alignments between human and *P. aurantium* orthologs, 11% of human minor introns are conserved as minor introns in *P. aurantium*, comparable to proportions shared between human and many plant species [56]; in alignments with *P. schiedti* the proportion of conserved human minor introns is closer to 2.5%, although this appears to largely be due to massive minor-to-major conversion of ancestral minor introns in *P. schiedti*, as 69% of the human minor introns in those alignments are paired with major introns in *P. schiedti*.

**Fig. 3.**
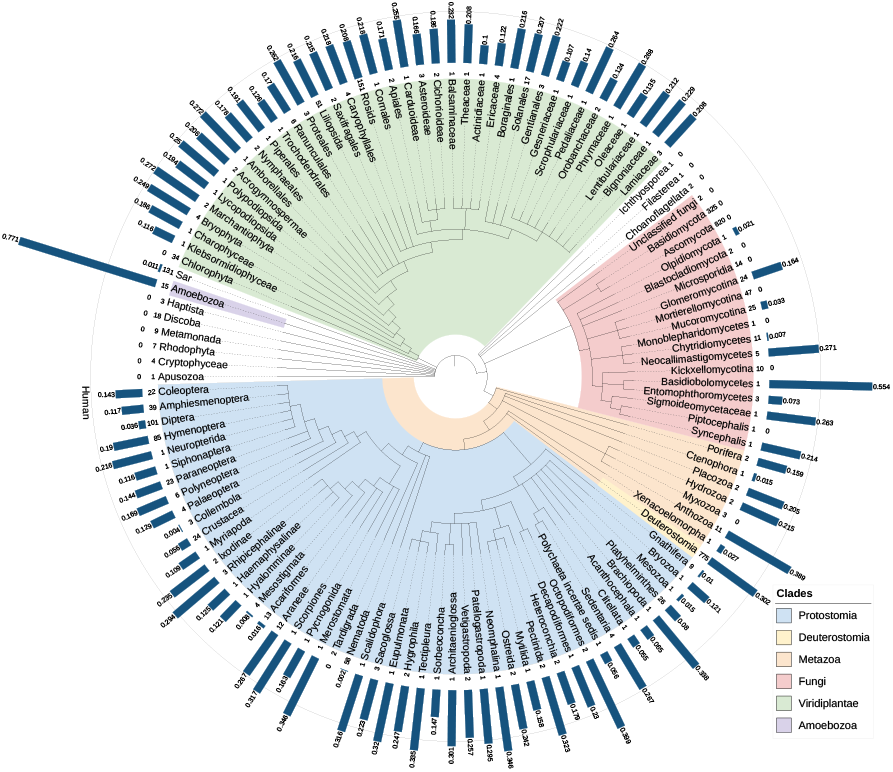
Average minor intron density (percentage of introns which are minor type; blue bars) in various eukaryotic clades. Integer numbers following clade names denote number of species represented. Outer circle indicates the minor intron density of the human genome. Taxonomic relationships based upon data from the NCBI Taxonomy Database [24].

As reported by Gentekaki et al. [28], the parasitic microbe *Blastocystis sp. subtype 1* within the stramenopiles contains hundreds of minor introns, although our pipeline identifies ∼45% fewer (n=253) than previously described. Interestingly, the *Blastocystis sp. subtype 1* minor introns we identify are highly enriched for the AT-AC subtype (77% or 196/253, where AT-AC introns are only ∼26% of all minor introns in human), and the classic minor intron bias away from phase 0 is inverted, with 49% (124/253) of the putative minor introns in *Blastocystis* being phase 0. *Blastocystis* also has the shortest average minor intron length in the data we analyzed at just under 42 bp (median 39 bp) (introns shorter than 30 bp were systematically excluded in all species).

Surprisingly, we find unusually high minor intron densities in a number of fungal species, a kindgom which until now was not known to contain significant numbers of minor introns. In particular, the Glomeromycete species *Rhizophagus irregularis* has a minor intron density comparable to that of humans (0.272%, n=205), and *Basidiobolus meristosporus*, in the Zoopagomycota, has one of the highest minor intron densities outside of the Amoebozoa (0.554%, n=249) (Fig. 4). We do not find any convincing support for minor introns in either of the two largest fungal groups, Ascomycota and Basidiomycota, which seem to have lost most if not all of the required minor snRNAs in the vast majority of species, as has been previously reported [48, 3].

**Fig. 4.**
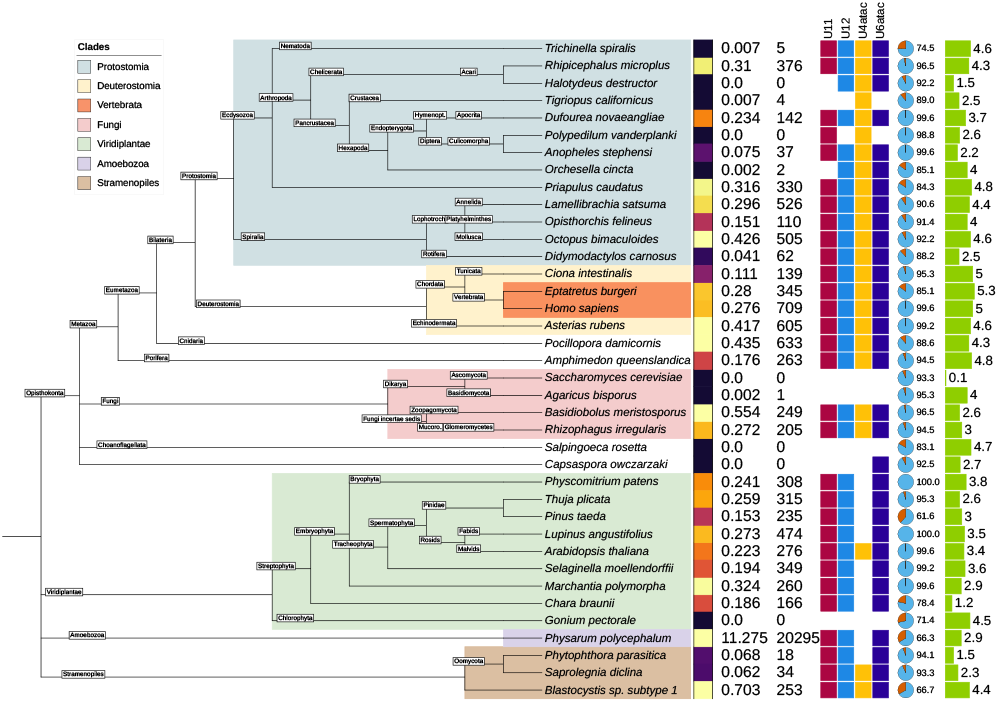
Minor intron densities and other metadata for selected species of interest. Graphical elements are as described in Fig. 1.

Our analysis confirms the presence of a small number of minor introns in the oomycete genus *Phytopthora* as reported by other groups [73, 3] (Fig. 4), and in addition we find that members of the stramenopile water mould genus *Saprolegnia* contain dozens of minor introns each. While any species with a very low number reported number of minor introns raises concerns about false positives, subsets of minor introns from each of these lineages have been found in conserved positions with minor introns in distantly-related species in our analyses, and minor snRNAs in each of the aforementioned genomes provides further evidence for the existence of bona fide minor introns in these species (Fig. 4). Interestingly, given its sister placement to the broadly minor-intron-poor nematode clade, the cactus worm *Priapulus caudatus* appears to be quite minor-intron rich (n=330, 0.316%), with substantial minor intron conservation to other metazoan lineages.

Within the protostomes, one of the two sister clades of bilateria, there are cases of relative minor intron enrichment in both arachnids (Arachnida) and molluscs (Mollusca) (Fig. 1, Fig. 4), as well as in the brachiopod species *Lingula anatina* and the horseshoe crab *Limulus polyphemus*. The order of ticks Ixodida, including *Ixodes scapularis, Dermacentor silvarum* and *Rhipicephalus*, has a much higher average minor intron density than other groups within Acari, which includes both mites and ticks and has seen substantial loss of minor introns in many of its lineages.

On the other side of the bilaterian tree, minor intron densities in deuterostomes are far more homogeneous. Vertebrates have consistently high minor intron densities (∼0.3%), with only a handful of exceptions in our data that are very likely due to incomplete or otherwise problematic annotations (for an example, see *Liparis tanakae* in Fig. 1, an individual species with dramatically lower minor intron densities than surrounding taxa, with a low `BUSCO` score and no indication of minor spliceosome loss). The remaining deeply-diverging clades within deuterostomes have minor intron densities comparable to vertebrates (the starfish *Asterias rubens* being on the high side of vertebrate densities, for example) with the exception of tunicates, which appear to have lost a significant fraction of their ancestral minor intron complement (and minor splicing apparatus, in the case of the highly-transformed species *Oikopleura dioica*).

In their seminal paper examining spliceosomal snRNAs in various eukaryotic lineages, Dávila López et al. [48] report a number of clades without some/any minor snRNAs. Based upon our larger dataset, it now seems clear that some of these groups do in fact have both minor introns and most if not all of the canonical minor snRNAs. These include the *Acropora* genus of coral, which has an average minor intron density higher than that of most vertebrates; within the fungal phylum Chytridiomycota the Chytridiomycete species *Spizellomyces punctatus* as well as a number of Neocallimastigomycetes including *Piromyces finnis* and *Neocallimastix californiae*; the genus of blood flukes *Schistosoma*; and all of the species of Streptophyta included in the previously-published analysis (see Fig. 1 in [48]). Notably, we also find minor introns (confirmed by comparative genomic methods) in the green algal species *Chara braunii* (n=166) and *Klebsormidium nitens* (n=110), representatives of a group which until now was thought to lack minor splicing entirely [48, 3, 73], as well as in the Glaucophyte alga *Cyanophora paradoxa* (n=77) (which may have transformed minor splicing machinery, as we find significant hits to only the U11 snRNA in that species).

#### 3.1.2 Minor intron depletion

Punctuated and dramatic loss of minor introns is a hallmark feature of the minor splicing landscape, and it remains an outstanding question why certain lineages undergo either partial or complete loss of their ancestral minor intron complements [90]. Previous work has delineated many groups that appear to lack either minor introns, minor splicing components or both [3, 73, 48], but the diversity and scope of more recently-available data motivated us to revisit this topic. In addition to the underlying data presented in Fig. 1, there are a number of cases of severe and/or complete minor intron loss that we highlight here. First, the amoebozoan *Acanthamoeba castellanii* has been found to contain both minor splicing apparatus as well as a limited number of minor-like intron sequences [73]. While it remains likely that this species contains a small number of minor introns based upon previous evidence, we do not find conservation of any of the twelve *Acanthamoeba* introns our pipeline classified as minor in either human or the more closely-related amoeobozoan *Protostelium aurantium*. This may not be particularly surprising, given the low absolute number of minor introns under consideration—between *Protostelium aurantium* and human, for example, ∼23% of *Protostelium* minor introns are conserved, and there are only two minor introns from *Acanthamoeba* in regions of good alignment with human orthologs. Furthermore, we do find a single shared minor intron position between *Acanthamoeba* and human when we disregard the local alignment quality and simply consider all introns in identical positions within aligned regions, which amounts to 20% of *Acanthamoeba* minor introns in such alignments.

Among clades with extreme but incomplete loss (a classic case in animals being Diptera), notable examples include the Acari (ticks and mites), bdelloid rotifers, and the springtail (Collembola) subclass of hexapods, We find no evidence at all for minor introns in the following taxa, many of which have not been reported before (those with citations corroborate earlier studies): tardigrades (e.g., *Hypsibius exemplaris*), Discoba (e.g., *Trypanosoma, Leishmania*) [48], Orchrophyta (stramenopiles), Alveolata (protists) [3, 48]. The Acari, in addition to an overall extreme reduction in minor introns within the clade generally, also contains a number of cases of apparent (by comparative genomic analysis) complete loss in the parasitic mite *Tropilaelaps mercedesae* (though minor introns are present in sister taxa) and the earth mite *Halotydeus destructor*. We also report two other novel cases of apparent complete minor intron loss outside of Acari. First, in the Dipteran clade Chironomidae, we find scant evidence of minor introns in *Clunio marinus, Polypedilum vanderplanki* and *Belgica antarctica*, all of which also appear to be missing between half and three-quarters of their minor snRNAs. Second, the copepod species of crustaceans *Tigriopus californicus* and *Eurytemora affinis* each lack both conserved minor introns and 75% of the minor snRNA set.

### 3.2 Minor introns have lower average conservation than major introns

A persistent result in the minor intron literature is that minor introns are more highly conserved than major introns (specifically, between animals and plants and even more specifically, between human and *Arabidopsis thaliana*) [4], although this assertion has been contradicted by at least one more recent analysis [56]. The claim that minor intron conservation exceeds major intron conservation largely rests upon the numbers of introns of both types found in identical positions within 133 alignments of orthologous human-*Arabidopsis* sequences, as reported in Table 1 of Basu et al. [4]. For major (U2-type) introns, they report 115 conserved as major in aligned ortholog pairs, and 1391 as either not present in one of the two orthologs or present as a minor intron; for minor (U12-type) introns, they find 20 conserved and 135 missing/converted. For each intron type, taking the number conserved and dividing by the total number of introns of that type present in the alignments results in conservation percentages of 7.6% 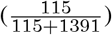 for major introns and 12.9% 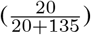 for minor introns (although the aforementioned values are not explicitly stated in the text), leading to the conclusion that minor introns are more highly conserved between human and *Arabidopsis* than are major introns. To the extent that we correctly understand their approach, however, we believe there may be a complication with this analysis.

**Table 1.**
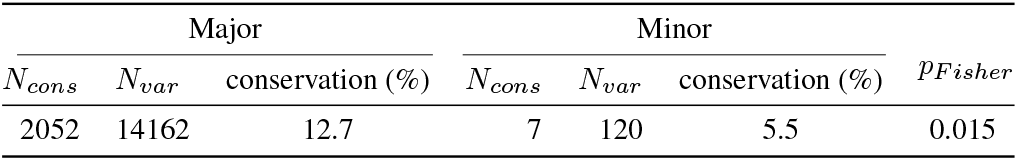
Comparison of major and minor intron conservation between human and *Arabidopsis thaliana. N*_*cons*_ indicates the number of introns of each type conserved as the same type in both human and *Arabidopsis. N*_*var*_ indicates the total number of introns (of both species) present in the alignments where the corresponding position in the opposing sequence either does not contain an intron, or contains an intron of the other type.

| Major |  |  | Minor |  |  | *PFisher* |
| --- | --- | --- | --- | --- | --- | --- |
| *N<sub>cons</sub>* | *N<sub>var</sub>* | conservation (%) | *N<sub>cons</sub>* | *N<sub>var</sub>* | conservation (%) |  |
| 2052 | 14162 | 12.7 | 7 | 120 | 5.5 | 0.015 |

Examining the ortholog pairs the authors provide in the supplementary data, it is evident that many of the same *Arabidopsis* sequences are present in multiple ortholog pairs, which suggests that a standard reciprocal-best-hit criteria for ortholog identification was not employed and that certain introns will be counted multiple times within the orthologous alignments. As many minor introns occur in larger paralogous gene families, this methodology could lead to artificial inflation of the calculated minor intron conservation, especially given the small absolute number of minor introns involved. To attempt to more thoroughly address the question of minor vs. major conservation, we identified orthologs in many different pairs of species across a range of evolutionary distances (see Methods), and calculated intron conservation using the same metric as above. Within more than 100 such comparisons between animals and plants (and more than 60 between animals and fungi), we find no cases where minor intron conservation exceeds major intron conservation (Fig. 5a).

**Fig. 5.**
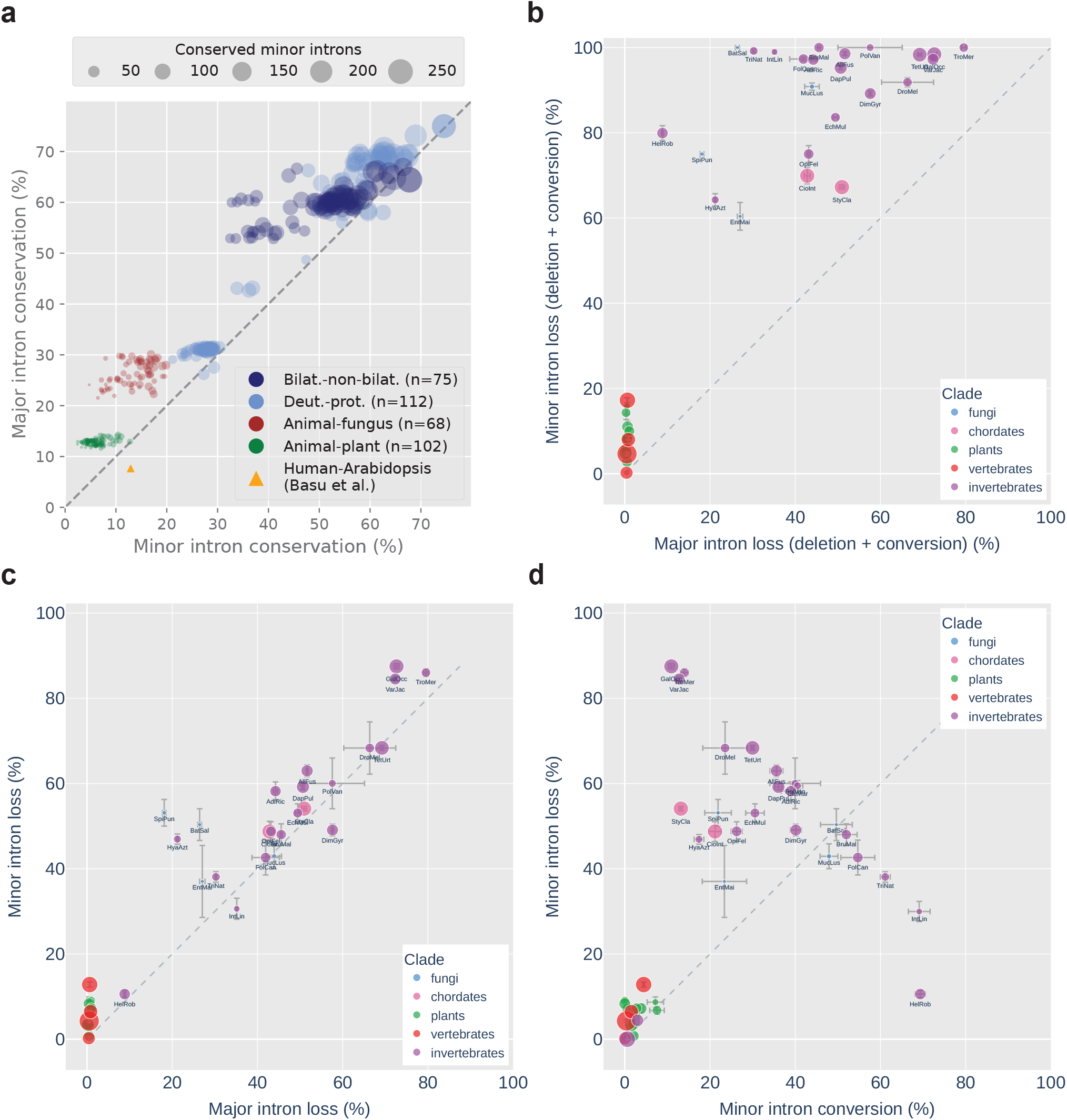
Conservation and loss of minor and major introns. (a) Comparison of major (y-axis) vs. minor (x-axis) intron conservation across hundreds of pairs of species. Bilat.-non-bilat.: bilaterian vs. non-bilaterian (animal); Deut.-prot.: deuterostome vs. protostome. The yellow triangle indicates levels of conservation of major and minor introns between *Homo sapiens* and *Arabidopsis thaliana* as reported by Basu et al. [4]. Size of markers indicates number of minor introns conserved between each pair. (b) Minor vs. major intron loss, where “loss” includes both sequence deletion and conversion to an intron of the other type. Bars indicate standard error of the mean for averaged values. Marker size represents relative minor intron density. Species abbreviations are as follow: AdiRic: *Adineta ricciae*, AllFus: *Allacma fusca*, BatSal: *Batrachochytrium salamandrivorans*, BruMal: *Brugia malayi*, CioInt: *Ciona intestinalis*, CluMar: *Clunio marinus*, DapPul: *Daphnia pulicaria*, DimGyr: *Dimorphilus gyrociliatus*, DroMel: *Drosophila melanogaster*, EchMul: *Echinococcus multilocularis*, EntMai: *Entomophaga maimaiga*, FolCan: *Folsomia candida*, GalOcc: *Galendromus occidentalis*, HelRob: *Helobdella robusta*, HyaAzt: *Hyalella azteca*, IntLin: *Intoshia linei*, MucLus: *Mucor lusitanicus*, OpiFel: *Opisthorchis felineus*, PolVan: *Polypedilum vanderplanki*, SpiPun: *Spizellomyces punctatus*, StyCla: *Styela clava*, TetUrt: *Tetranychus urticae*, TriNat: *Trichinella nativa*, TroMer: *Tropilaelaps mercedesae*, VarJac: *Varroa jacobsoni*. (c) Minor vs. major intron loss, where “loss” represents actual deletion of the intron sequence. Other elements as in Fig. 5b. (d) Minor intron loss vs. conversion, where “loss” represents actual deletion of the intron sequence. Other elements as in Fig. 5b.

Furthermore, we observe only a handful of cases where minor intron conservation marginally exceeds major intron conservation in alignments of more closely-related species (∼3% greater between the starfish *Asterias rubens* and the stony coral *Orbicella faveolata*, for example). In the specific case of human-*Arabidopsis* considered by Basu et al., our data show minor intron conservation to be around half that of major intron conservation (Table 1). Thus, in the final analysis we find no compelling support for the idea that minor introns are in general more conserved than major introns and in fact, the opposite seems to be true in the vast majority of cases.

### 3.3 Minor intron loss vs. conversion

When an ancestral minor intron ceases to be a minor intron, it is thought to happen primarily in one of two ways: the entire intron sequence could be lost via, for example, reverse transcriptase-mediated reinsertion of spliced mRNA [44, 17, 90, 33], or the intron could undergo sequence changes sufficient to allow it to be recognized instead by the major spliceosome [11, 21, 20, 26]. From first-principles arguments based on the greater information content of the minor intron motifs [11, 10, 21] along with empirical analyses [44], it is assumed that intron conversion proceeds almost universally unidirectionally from minor to major. Previous work has also shown that the paradigm of full intron loss (sequence deletion) appears to dominate over conversion in minor introns [44]; we wondered whether any exceptions to this general pattern might exist.

First, we assembled a manually-curated sample of species with significant/complete minor intron loss, along with a number of species with much higher minor intron conservation for comparison. For each selected species, we chose an additional species to compare against as well as a species to serve as an outgroup, and then identified orthologs between all members of the group to allow us to identify ancestral major/minor introns (see Methods for details) and estimate fractions of each intron type retained. Considering loss to include both sequence deletion as well as type conversion (which we assume to be unidirectional from minor to major, as discussed above), Fig. 5b shows that minor intron loss is more pronounced than major intron loss in the species we examined (also shown more generally in Fig. 5a).

We can, however, also decompose the phenomena contributing to the higher degree of loss of minor introns and ask whether the contribution of sequence deletion specifically, for example, differs between the two types. Somewhat surprisingly, we find that this form of intron loss is very similar between the two types of introns in species which have lost significant fractions of their minor introns (Fig. 5c). Because the selected species were chosen based upon putative loss of minor introns and the sample size is low, it is difficult to interpret the apparent bias toward minor intron deletion in the vertebrates and plants in Fig. 5c. For the other species, however, this data suggests that there is not a particular selective pressure toward removing minor intron sequences themselves—at least not any more than there is pressure to remove intron sequences generally—in instances of pronounced minor intron upheaval.

We can also look at the other side of the minor intron loss coin, namely conversion from minor to major type. Here, we find that in many instances loss via deletion does indeed outstrip conversion (as reported by [44]), sometimes dramatically so, but there are interesting exceptions. The leech *Helobdella robusta* (HelRob), for example, which seems to have retained a large fraction of its ancestral major introns, has lost ∼80% of its minor introns primarily through conversion to major-type (Fig. 5d). By contrast, the annelid worm *Dimorphilus gyrociliatus* (DimGyr), found in a clade (Polychaeta) sister to *Helobdella robusta*, has undergone a seemingly independent loss of minor introns of similar proportion to *Helobdella* under a very different modality, with loss (deletion) outweighing conversion (Fig. 5d). It is unclear what forces are responsible for the relative contributions of each mechanism; in *Helobdella*, the major intron sequences are slightly more degenerate at the 5^*′*^SS end than in e.g., human, which might lower the barrier to entry for would-be minor-to-major converts. This, however, is mere speculation and more work is needed to better characterize these dynamics. It should be noted that under the current analysis we cannot differentiate between losses, and conversions followed by subsequent loss. Our conversion estimates, therefore, should be taken as lower bounds.

### 3.4 Positional biases of major and minor introns

It has been known for many years that introns often exhibit a 5^*′*^ bias in their positions within transcripts [55, 45, 74]. This can be explained in large part due to biased intron loss: because a primary mechanism of intron loss (and, to a more limited extent, gain) is thought to occur via the reverse-transcriptase mediated (and 3^*′*^-biased) insertion of spliced mRNA [70, 72, 19], over time such a process would tend to result in higher concentrations of introns closer to the 5^*′*^ end of transcripts.

Less attention has been paid to the positional biases of minor introns specifically, although at least one study [4] found that minor introns appear to be especially over-represented in the 5^*′*^ portions of transcripts in both human and *Arabidopsis thaliana*. We were curious to see whether the same patterns were present in our own data and whether they generalized beyond the two species so far examined.

We selected two sets of species to highlight—for the first, we chose lineages with substantial numbers of minor introns from a variety of groups; for the second, we picked species with significant inferred amounts of minor intron loss to investigate whether any 5^*′*^ bias might be more extreme in the remaining minor introns. In our analysis, we confirm the 5^*′*^ bias as previously described [4] in *Arabidopsis thaliana* (Fig. 6a), although we do not find the same difference as shown in the earlier study between major and minor intron positions in human.

**Fig. 6.**
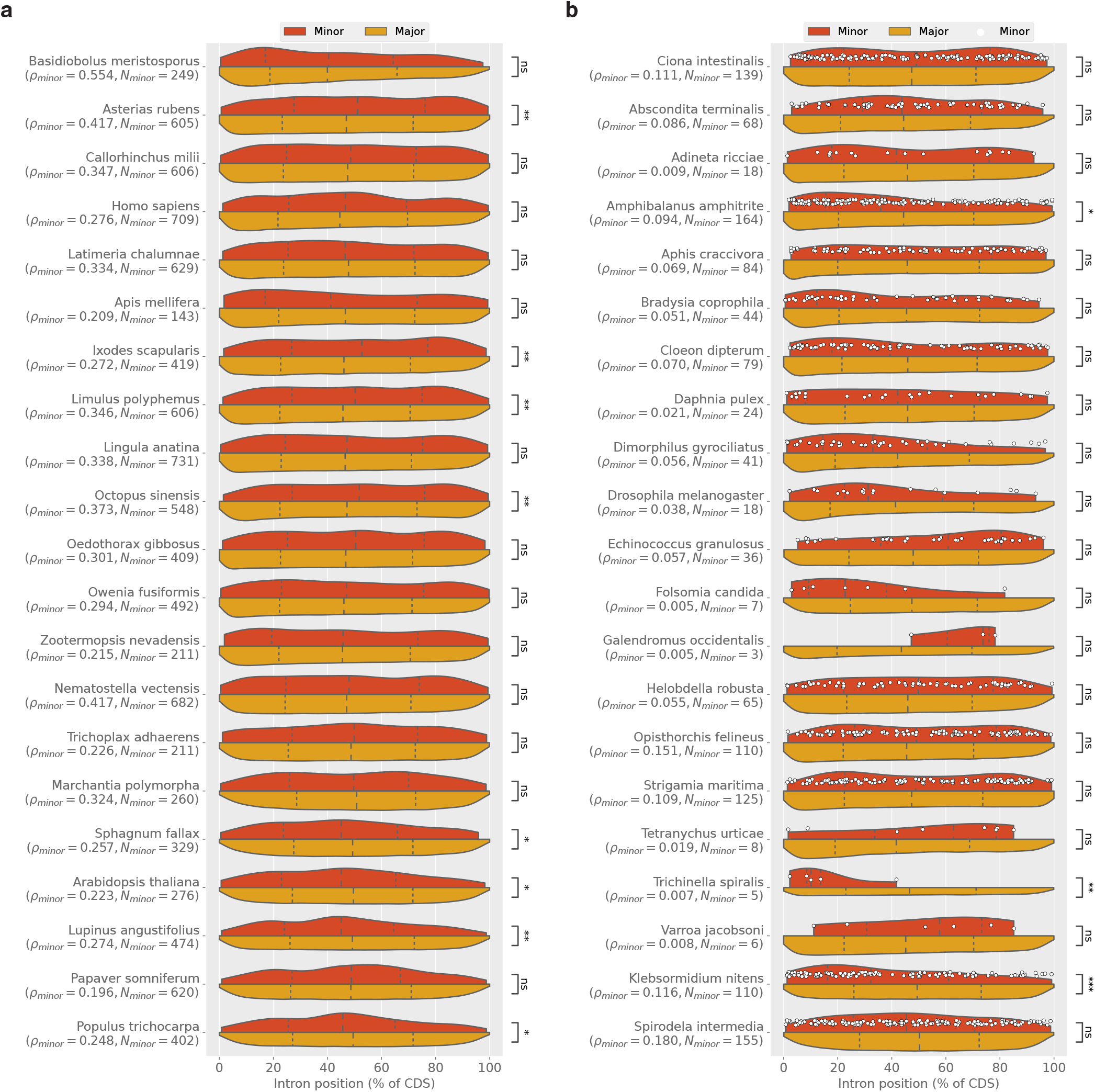
Intron position distributions for major (red) and minor (yellow) introns in selected species. (a) Species enriched in minor introns. (b) Species with significant inferred minor intron loss; white dots represent individual minor introns. For both plots: Dashed lines represent the first, second and third quartiles of each distribution. Statistically significant differences between minor and major introns are indicated with asterisks (two-tailed Mann-Whitney U test; * *p* < 0.05; ** *p* < 0.001; *** *p* < 0.0001; `ns`=not significant). Note that in some cases of significant differences between the two intron types, e.g., within animals, the set with greater 5^*′*^ bias is the *major* introns.

More broadly, our results point to a less-clear picture than earlier work might suggest—while we do find a number of cases in animals where minor introns are more 5^*′*^-biased than major introns (Fig. 6b, *Amphibalanus amphitrite* and *Trichinella spiralis*), the pattern is not broadly significant and is occasionally reversed (e.g., *Ixodes scapularis*), albeit in animal species with less-dramatic minor intron loss (Fig. 6a). Within plants, however, there does seem to be a clearer pattern, with a much higher fraction of plants species in both groups displaying a strong 5^*′*^ bias in their minor introns. To determine how widespread this pattern of greater relative 5^*′*^ bias in minor introns is, we searched our entire dataset for species with a) significant differences in minor intron occurrence between the 5^*′*^ and 3^*′*^ halves of trancripts (as assessed by a two-tailed exact binomial test, where presence in the 5^*′*^ half of a transcript was considered a success, mirroring the approach of ref. [4]), b) significant differences between major and minor positions as determined by a two-tailed Mann-Whitney U test and c) median minor intron positions more 5^*′*^ biased than median major intron positions. Among such species, plants are highly over-represented (Table 2, *p* = 8.9 *×* 10^−68^ by a Fisher’s exact test).

**Table 2.** Proportions of species in various groups with statistically significant 5*′* bias (*N*_5*′MIB*_) of minor intron positions within transcripts.

| Clade | $N_{total}$ | $N_{5'MIB}$ | % |
| --- | --- | --- | --- |
| Streptophyta | 290 | 112 | 38.6 |
| Fungi | 63 | 2 | 3.2 |
| Metazoa | 1204 | 21 | 1.7 |
| Stramenopiles | 16 | 0 | 0.0 |
| Evosea | 4 | 0 | 0.0 |
| Discosea | 1 | 0 | 0.0 |

It is possible that this pattern, taken together with the higher degree of stability of minor intron densities in plants, reflects an ancient loss of minor introns in the plant ancestor, the signature of which is now shared broadly among extant species. It might also suggest a unique and/or more consistent paradigm for minor intron loss in plants, distinct from the haphazard process seemingly at work within other parts of the eukaryotic tree where losses have occurred both more recently and more frequently.

### 3.5 Phase biases of minor introns

Spliceosomal introns can occur at one of three positions relative to protein-coding sequence: between codons (phase 0), after the first nucleotide of a codon (phase 1) or after the second (phase 2). In most species, major introns display a bias toward phase 0^2^ [59, 47] (Fig. 7a), while minor introns are biased away from phase 0 [11, 56] (Fig. 7b).

**Fig. 7.**
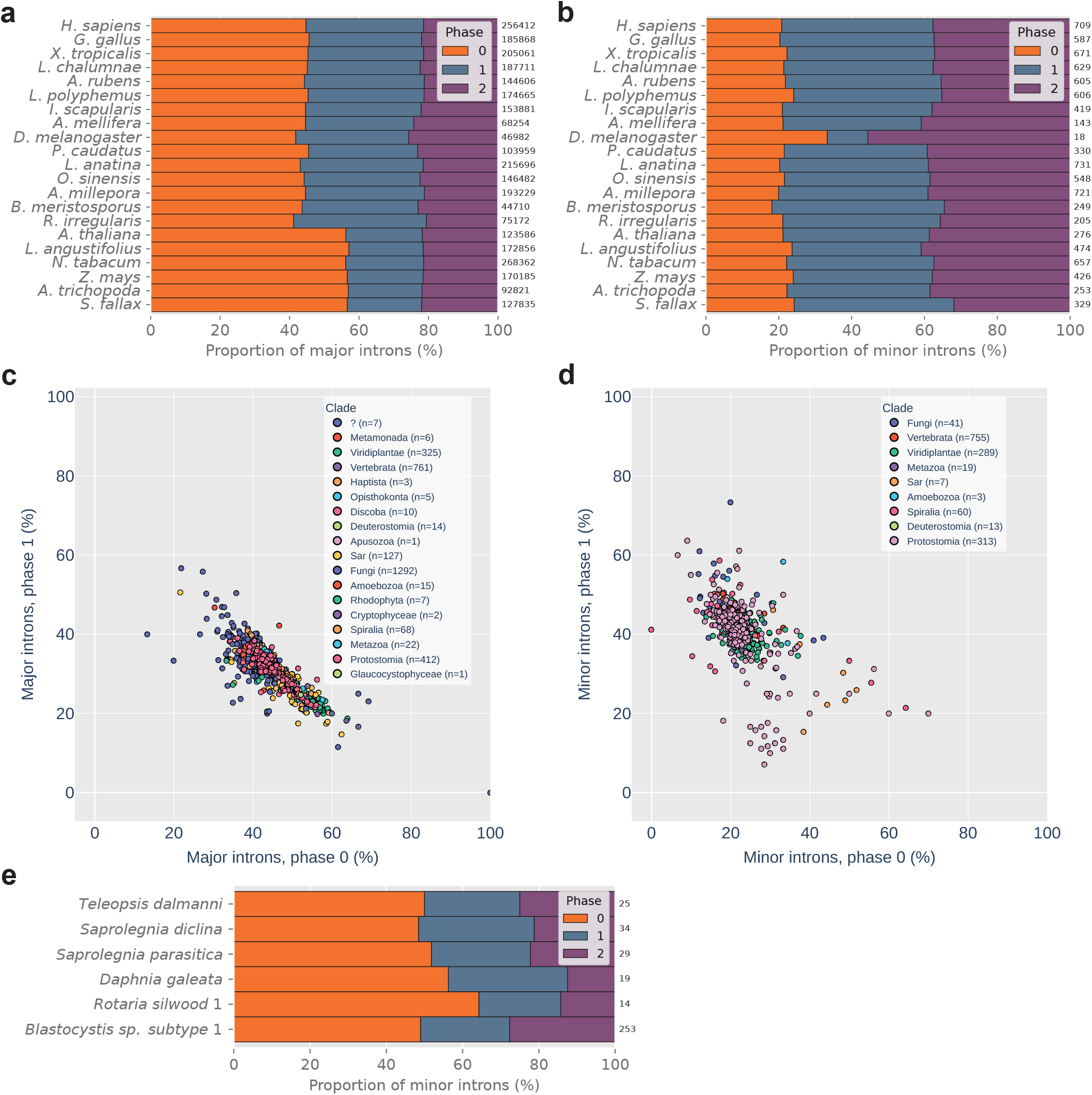
Minor and major intron phase biases. (a) and (b): Phase distributions of major and minor introns, respectively, in various species. Numbers at the ends of bars represent the total number of constituent introns. (c) Proportions of phase 1 (y-axis) vs. phase 0 (x-axis) for major introns in species with minor introns. Correlation of phase 0 to phase 1: *ρ*_*s*_ = −0.81, *p* = 0.0. (d) Proportions of phase 1 (y-axis) vs. phase 0 (x-axis) of minor introns in various species. Correlation of phase 0 to phase 1: *ρ*_*s*_ = −0.48, *p* = 2.14 *×* 10^−87^ (e) Unusually high proportions of phase 0 minor introns in certain species. Numbers at the end of each bar represent the total number of constituent introns. Proportions of phase 0 minor introns for all species are significantly different from expected values derived from the proportion of phase 0 minor introns in human (phase 0 vs. sum of other phases, Boschloo’s exact test *p <* 0.05).

It remains an unsettled issue why minor introns are biased in this way—one theory proposed by Moyer et al. [56] suggests that such bias could arise from preferential conversion of phase 0 minor introns to major type, which would over time lead to the observed pattern. Here, we wanted to make use of the size of our dataset to better characterize the diversity of intron phase more broadly, and identify any exceptions to the general rule. As shown in Fig. 7c, the phase distributions of major introns are fairly tightly grouped. On average, for major introns in minor-intron-containing species, phase 0 makes up 47%, phase 1 30% and phase 2 23%. In addition, the proportions of phase 0 and phase 1 introns are quite highly correlated (see caption of Fig. 7c). Minor introns, on the other hand, are less consistent in their phase distribution and have a lower phase 0 to phase 1 correlation, although the majority cluster relatively tightly around the average value of phase 0, 22% (Fig. 7d).

It is intriguing that a small number of species appear to have much higher fractions of phase 0 minor introns (Figures 7d and 7e). What’s more, these species (with the notable exception of *Blastocystis sp. subtype 1*, addressed below) all have very low absolute numbers of minor introns (Fig. 7e). While these data are not necessarily incompatible with the conversion paradigm mentioned above (which might predict minor introns in species with pronounced loss to show especially strong bias away from phase 0, although with such small numbers of remaining minor introns it may simply be that stochasticity dominates), it at least invites further investigation into the forces underlying the phase biases in minor introns generally.

The species *Blastocystis sp. subtype 1* is similar to the other unusual cases mentioned in its reduced minor intron bias away from phase 0, but is remarkable for the number of minor introns involved (n=253). It is also interesting that its minor intron phase distribution is almost identical to the distribution of phases in its major introns (not shown). While this raises the possibility that the minor introns in *Blastocystis sp. subtype 1* are false-positives, the fact that we find a) all four minor snRNAs in the genome, b) a (small but non-zero) number of its minor introns conserved in *Lingula anatina* (not shown) and c) putative minor introns in a closely-related species (*Blastocystis hominis*) provides evidence that they are likely real. Assuming they are bona fide minor introns, another possible explanation for the phase 0 enrichment could be that they have been more recently gained, and (under the conversion hypothesis) have not yet had time to develop the phase bias present in older minor intron sets. More thorough comparative genomics work within the clade after additional species become available would help to clarify the evolutionary picture.

### 3.6 Non-canonical minor intron splice boundaries

The vast majority (*>* 98.5%) of major introns in most eukaryotic genomes begin with the dinucleotide pair GT, and end with the pair AG [12, 13, 56, 80], with an additional much smaller contingent of GC-AG introns present in many genomes. When minor introns were first discovered, they were initially characterized largely by their distinct AT-AC termini [30, 34]. However, it was subsequently discovered that in fact the majority of minor introns in most species share the same terminal boundaries as major introns [21, 11], although the AT-AC subtype may constitute a more significant fraction of minor introns in certain species [90, 65, 56, 1, 3]. Over time, additional non-canonical (i.e., not GT-AG, GC-AG or AT-AC) subtypes of minor introns have been identified in various organisms [61, 1, 56, 80, 44], but these analyses have been limited to species for which there were available minor intron annotations which until now were quite limited.

Because non-canonical introns do not (by definition) look like normal introns, it can be difficult to differentiate between biological insights and annotation errors when examining eukaryotic diversity at scale. For example, a recent report on non-canonical introns in diverse species described significant enrichment of CT-AC introns in fungi [25]. However, and as addressed briefly in the paper itself, CT-AC boundaries happen to be the exact reverse-complement of the canonical GT-AG boundaries. Additional sequence features of these introns, such as a high occurrence of C 5 nt upstream of the 3^*′*^SS (which would perfectly match the hallmark +5G were the intron on the other strand) and an enrichment in +1C after the 3^*′*^SS (corresponding to the canonical -1G at the 5^*′*^SS on the other strand) make it very likely in our estimation that such introns are in fact incorrectly annotated due to some combination of technical errors and antisense transcripts. To combat issues of this sort, we first performed multiple within-kingdom alignments of various animal and plant species with high relative levels of annotated non-canonical minor intron boundaries. Conserved introns were then clustered across many different alignments to form conserved intron sets, which were then filtered to include only minor introns in sets where at least two minor introns were found (see Methods for details). These sets of introns are much less likely to contain spurious intron sequences, although they also may not fully represent more recent or lineage-specific boundary changes and they do not include introns from every species in our collected data.

Our results in animals (Fig. 8a) and plants (Fig. 8b) are largely consistent with previous data on non-canonical minor introns [61, 80, 44], with only small differences in the rank-order within each set. The set of plant non-canonical minor intron termini is both less-diverse and more lopsided than the animal set; while the most common non-canonical termini is AT-AA in both kingdoms, almost 75% of all non-canonical minor introns we identify in plants are of the AT-AA subtype, versus less than half that proportion for the same subtype in animals. Interestingly, the second most common non-canonical termini in animals, AT-AT, is almost entirely absent in plants.

**Fig. 8.**
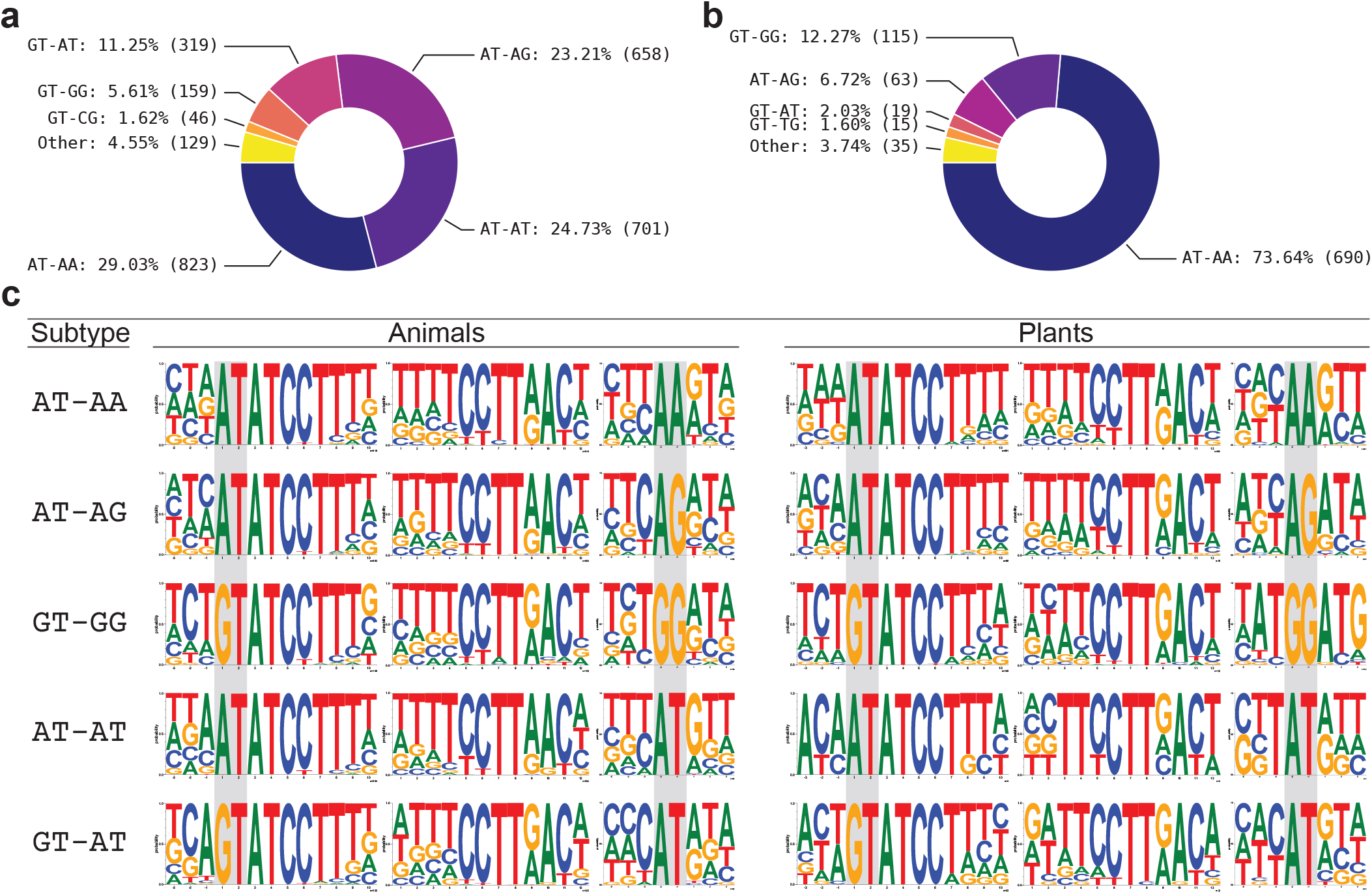
Non-canonical minor intron motifs in animals and plants. (a) and (b): Non-canonical intron termini found in conserved minor introns in animals and plants, respectively. (c) Sequence motifs of the 5*′*SS, BPS and 3*′*SS regions of non-canonical minor introns in animals and plants. The terminal dinucleotide pairs are highlighted in gray.

As can be seen in Table 3, the majority of non-canonical termini differ by a single nucleotide from a canonical terminus; only GT-TA, GT-CA, AT-GA, AT-CG, CT-AT, and AT-GT in animals and AT-TT, AT-CA, AT-CG, AT-GA, and AT-GT in plants differ by more than one nucleotide, and each are only a tiny minority of the total non-canonical set. Additionally, there are small differences between the consensus sequences outside of the terminal dinucleotides between the different subtypes of minor introns (Fig. 8c), and also within the same subtype between animals and plants. The most prominent examples of the latter are in the following subtypes: GT-GG (AT motif immediately preceeding the 3^*′*^SS, and ATG motif immediately following it in plants), AT-AG (−1C from the 3^*′*^SS in animals) and GT-AT (−1A in animals).

**Table 3.** Non-canonical minor intron termini in animals and plants. Termini with only a single occurrence are excluded.

| Animals |  |  | Plants |  |  |
| --- | --- | --- | --- | --- | --- |
| Termini | % | Count | Termini | % | Count |
| AT-AA | 29 | 823 | AT-AA | 73.6 | 690 |
| AT-AT | 24.7 | 701 | GT-GG | 12.3 | 115 |
| AT-AG | 23.2 | 658 | AT-AG | 6.7 | 63 |
| GT-AT | 11.3 | 319 | GT-AT | 2 | 19 |
| GT-GG | 5.6 | 159 | GT-TG | 1.6 | 15 |
| GT-CG | 1.6 | 46 | TT-AG | 1.3 | 12 |
| GT-TG | 1 | 29 | AT-CA | 0.6 | 6 |
| CT-AC | 0.8 | 24 | GT-CG | 0.5 | 5 |
| GG-AG | 0.7 | 21 | AT-AT | 0.4 | 4 |
| GA-AG | 0.5 | 14 | AT-TT | 0.2 | 2 |
| TT-AG | 0.4 | 10 |  |  |  |
| GT-TA | 0.1 | 3 |  |  |  |
| GT-CA | 0.1 | 3 |  |  |  |
| AT-GA | 0.1 | 3 |  |  |  |
| AT-GC | 0.1 | 3 |  |  |  |
| CT-AG | 0.1 | 2 |  |  |  |
| AT-CG | 0.1 | 2 |  |  |  |
| CT-AT | 0.1 | 2 |  |  |  |
| AT-CC | 0.1 | 2 |  |  |  |
| AT-GT | 0.1 | 2 |  |  |  |

### 3.7 Minor intron-containing genes are longer and more intron-rich than genes with major introns only

Across the eukaryotic tree, genomes can vary widely in the number of introns contained in an average gene [72]. Some vertebrate genes have dozens or even hundreds of introns (e.g., the gene titin in human), whereas most genes of the yeast *Saccharomyces cerevisiae* lack introns entirely. Given the fact that minor introns appear to be arranged non-randomly throughout genomes where they are found [56, 43, 80, 1] a natural question to ask is to what extent and in what ways are minor intron-containing genes (MIGs) different than those without minor introns? As far as we are aware, while various aspects of this question have been addressed by different groups [4, 11, 35], relatively little attention has been paid to possible differences in a number of basic gene attributes, namely gene length (excluding introns) and number of introns per unit coding sequence or “genic intron density” (a coinage we will use here to distinguish from our more frequent usage in this paper of “intron density” to describe some number of introns in terms of their relative share of the total introns in the genome).

Strikingly, when we compare the genic intron density of MIGs to all other genes in species with minor introns, we find that MIGs are universally more intron-dense on average than non-MIGs (Fig. 9a). Furthermore, it appears that average MIG lengths (excluding intron sequences) are longer than other genes in the vast majority of species with minor introns (Fig. 9b). While there are a number of cases where the median non-MIG gene length exceeds the median MIG gene length, none of those differences are statistically significant (Mann-Whitney U test, *p* ≥ 0.05). An in-depth analysis of this qualitative finding is beyond the scope of the current paper, but it seems an underappreciated difference between the two intron types and would benefit from further investigation.

**Fig. 9.**
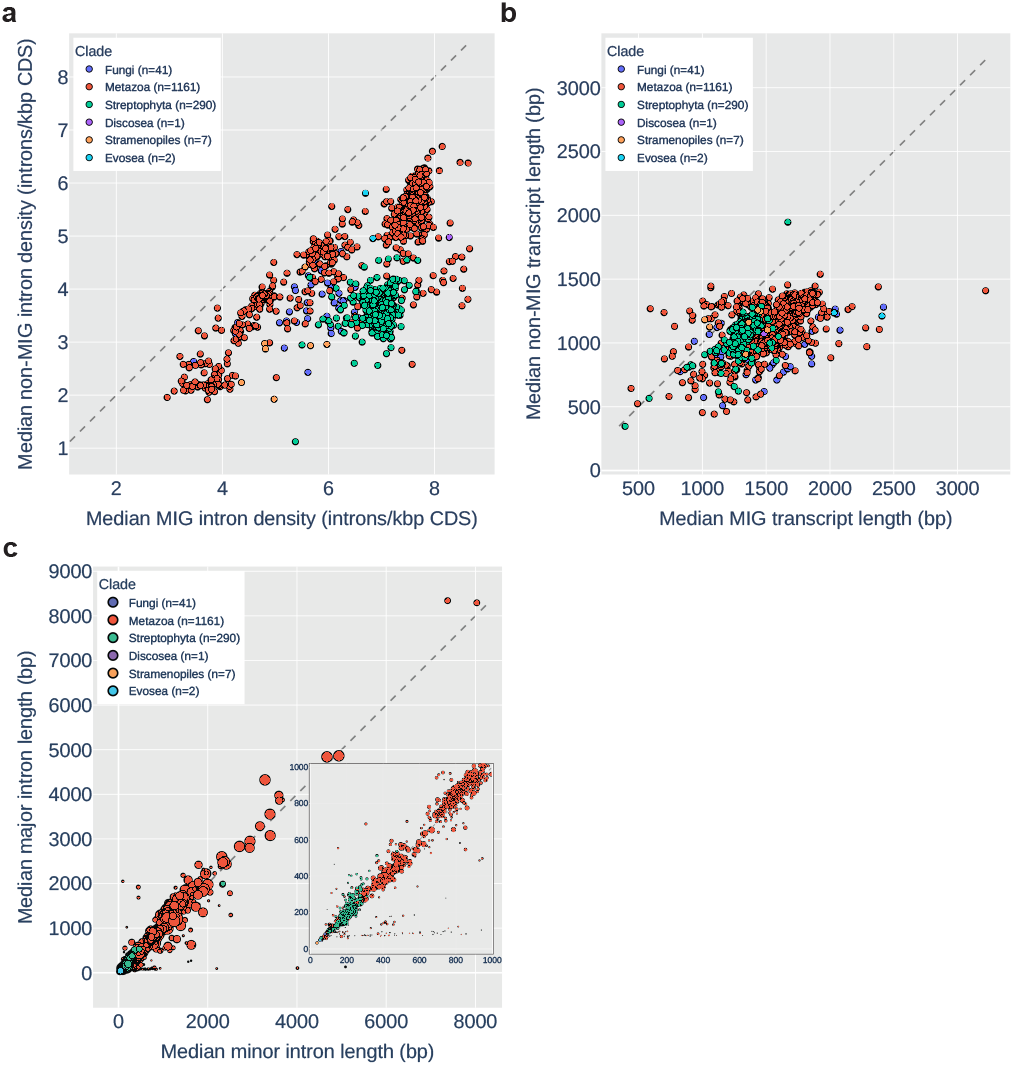
Features of MIGs and minor introns. (a) and (b): Median genic intron density (introns/kbp coding sequence) and gene length (sum of CDS), respectively, for major-intron-only genes (y-axis) vs. minor intron-containing genes (x-axis). (c) Median major intron length (y-axis) vs. median minor intron length (x-axis) for all species with high-confidence minor introns. Size of markers indicates number of minor introns in the genome. Inset: The subset of the same data with length ≤ 1000 bp.

### 3.8 Comparison of minor and major intron lengths

While a number of studies have compared the length distributions of the different intron types in a limited assortment of genomes [43, 91, 56], without a large set of minor intron containing species to compare within it has been difficult to gauge the extent to which minor intron lengths might differ from major intron lengths. With the comprehensive minor intron data we have collected, we were able to ask a very basic question: what is the general relationship between average major and minor intron lengths? At a high level, the answer appears to be that major and minor intron lengths are roughly linearly correlated (Fig. 9c)—species with longer average major intron length tend to also have longer average minor intron length (Spearman’s *ρ* = 0.625 for median values, *p* = 5.08 *×* 10^−16^). One interesting aspect of the data in Fig. 9c is shown more clearly in the inset plot (which is simply the subset of the data in the main plot with length ≤ 1000 bp): certain species have significant minor intron loss (small markers) and large differences between average minor and major intron lengths. It should be noted that the set of species in that region is enriched for *Drosophila* (a taxonomically over-represented genus in the sequence databases), but includes many additional insect species as well.

Although it is not clear what immediate conclusions can be drawn from this data, some additional questions are raised: Were shorter minor introns especially selected against in these lineages, such that the remaining minor introns are disproportionately long? What is driving variation within, for example, *Drosophila* such that in some species the difference between minor and major is relatively modest (*Drosophila busckii*, major=65 bp and minor=189 bp) and in others, it’s much more stark (*Drosophila biarmipes*, major=77 bp and minor=677 bp)? It should be noted as well that for *Drosophila* specifically, almost all of the minor introns are conserved within the genus, so the previous example is made more interesting because 100% of the *D. busckii* minor introns are shared with *D. biarmipes*, yet are far longer in the latter than the former. It did occur to us to check whether minor introns in these outlier species happen to be (for whatever reason) in genes with longer-than-average intron size, and although we have not done so systematically we have checked a number of more extreme cases and have found the same pattern recapitulated between minor and major introns of the same genes. For example, in the black soldier fly *Hermetia illucens*, the median minor intron length is 4019 bp and the median major intron length is only 105 bp. Comparing minor to major within only the minor intron-containing genes changes things, but not qualitatively—the median major intron length becomes 399.5 bp, but the difference between minor and major is still significant (*p* = 0.0025 by one-tailed Mann-Whitney U test under the alternative hypothesis that minor intron lengths are longer).

### 3.9 Reconstruction of ancestral minor intron densities

In an attempt to quantify some of the evolutionary dynamics leading to the variegated pattern of minor intron densities we see in extant lineages, we sought to estimate minor intron densities for certain ancestral nodes throughout the eukaryotic tree (see Methods). For each selected node, we identified pairs of species for which the node is the most recent common ancestor and, in combination with an outgroup species, performed three-way protein-level alignments to allow us to define intron states for each species within the alignments. Then, using the procedure described in [69], we calculated the number of minor and major introns estimated to have been present in the aligned regions in the ancestral genome, and repeated this process using many different combinations of species for each node to derive average values across all such comparisons. Because the absolute number of introns present in the aligned regions in the ancestor is not a particularly easy value to interpret, for reconstructions within a given kingdom we normalized the ancestral density of each intron type by a chosen reference species from that kingdom present in every alignment (see Methods for details). The reference species for animals, fungi and plants were *Homo sapiens* (minor intron density 0.276%), *Rhizophagus irregularis* (minor intron density 0.272%) and *Lupinus angustifolius* (minor intron density 0.273%), respectively. Fig. 10 shows both distributions of minor intron densities in constituent species from each terminal clade (violin plots), as well as estimates of ancestral minor intron densities at various nodes (colored boxes) as fractions of the density of minor introns in the aligned regions of the reference species (i.e., ancestral densities > 1 indicate minor intron enrichment relative to the reference species, and ancestral densities *<*1 indicate reduction).

**Fig. 10.**
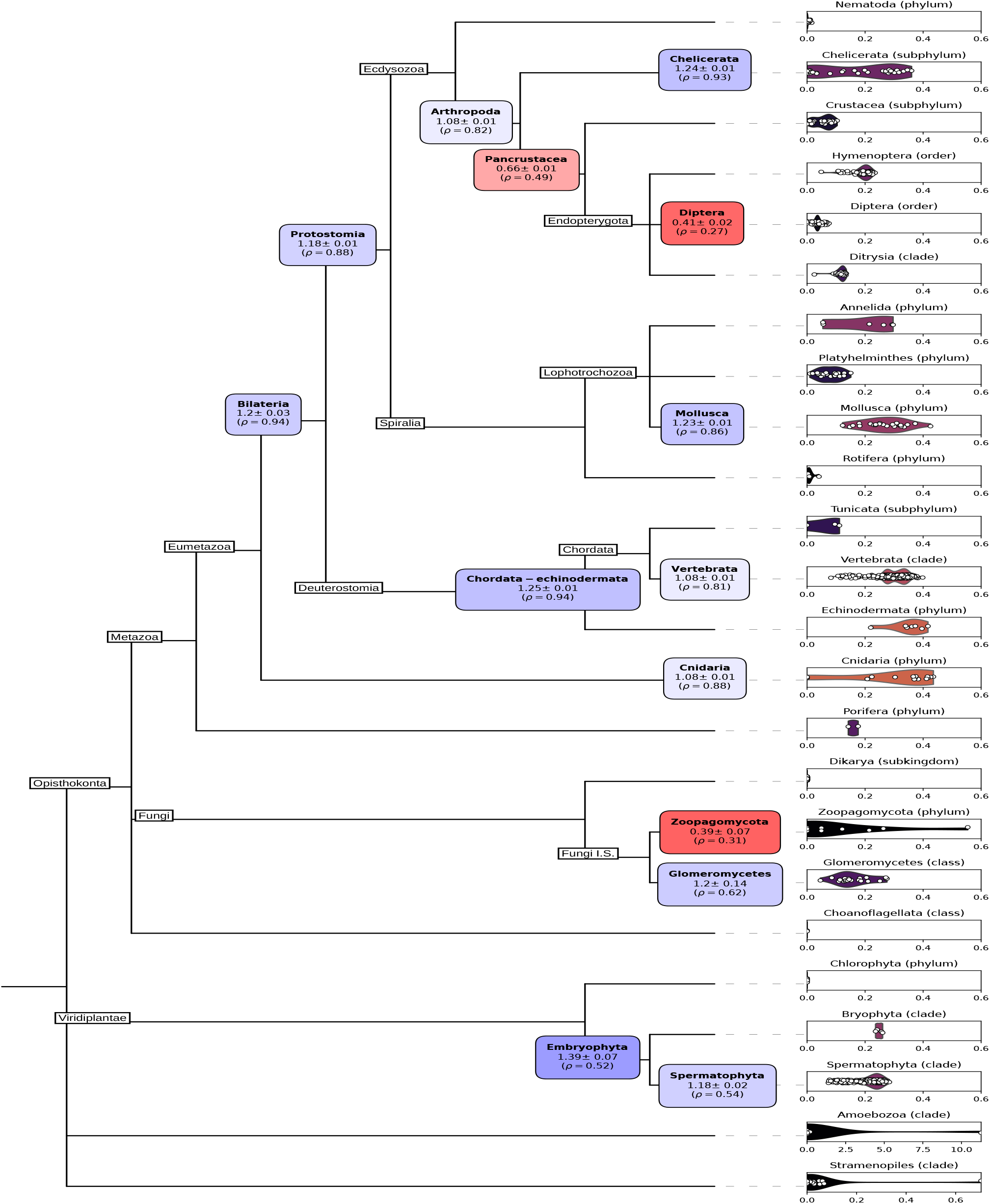
Minor intron density distributions in selected clades, and ancestral reconstructions of minor intron densities at selected nodes. Ancestral density node label color indicates relative enrichment (blue) or reduction (red) relative to the reference species in the alignments. For animals, the reference is *Homo sapiens*; for plants the reference is *Lupinus angustifolius;* for fungi the reference is *Rhizophagus irregularis*.

As shown in Fig. 10, ancestral minor intron densities were, in large part, modestly higher than minor intron densities in the relatively minor-intron-rich reference species, with the exception of a number of episodes of pronounced loss in the ancestors of Diptera, Pancrustacea and Zoopagomycota. The apparent enrichment of minor introns in the ancestor of Chelicerata is interesting, as it suggests there may have been some amount of minor intron gain along that branch from the arthropod ancestor. This result needs to be qualified, however, by noting that in that region of the tree we were constrained by lack of available data to using only *Limulus polyphemus* for one of the two ingroup species, as well as the fact that in any given reconstruction, the calculated intron density is limited to the genes involved in the reconstruction. With similar caveats, the low inferred ancestral minor intron density of Zoopagomycota is notable as that group contains *Basidiobolus meristosporus*, which has the highest minor intron density so far discovered in fungi (0.554%). Overall, these results paint a picture of ancestral minor intron complements as generally analogous to those of minor intron rich extant species, and highlight the quixotic nature of minor intron loss dynamics throughout eukaryotic diversity. It would be interesting to have these results expanded upon once phylogenetic uncertainty has been reduced throughout the tree and even more diverse genomes are available for analysis.

### 3.10 Unprecedented minor intron density in the fungus *Rhizophagus irregularis*

In our broad survey of eukaryotic species, we found a large number of putative minor introns in the mycorrhizal fungus *Rhizophagus irregularis*, a member of the Glomeromycota group of fungi. There is clear correspondence between minor-versus-major spliceosomal sequence characteristics in the two primary differentiating parts of the introns, namely the 5^*′*^ splice site and the 3^*′*^ branchpoint structure (Fig. 11a), and consensus sequence features closely follow those previously found in animals and plants (Fig. 11b). A subset of minor introns were found at conserved gene positions with minor introns in other fungi, animals and plants (Fig. 11c,d), further increasing our confidence that these introns represent bona fide minor spliceosomal introns. Searches of the genome provided further evidence for presence of many minor spliceosome-specific proteins as well as all four minor spliceosome-specific non-coding RNAs (U11, U12, U4atac, and U6atac) (Fig. 11e,f). As found previously in animals and plants, we found a distinctive distribution of intron phase (position at which introns interrupt the coding codon series, whether between codons (phase 0) or after the first or second nucleotide of a codon (phase 1 and 2, respectively): whereas major introns typically show the pattern (ph0 > ph1 > ph2), minor introns in *R. irregularis* followed the minor pattern in animals and plants (ph1 > ph2 > ph0; [43, 56]) (Fig. 11g). In total, we predict that 199 introns in *R. irregularis* are minor-type (0.275% of 72,285 annotated introns), orders of magnitude higher than for other fungal species previously reported to contain minor introns (∼4 in *Rhizopus oryzae* and ∼20 in *Phycomyces blakesleeanus*) [3].

**Fig. 11.**
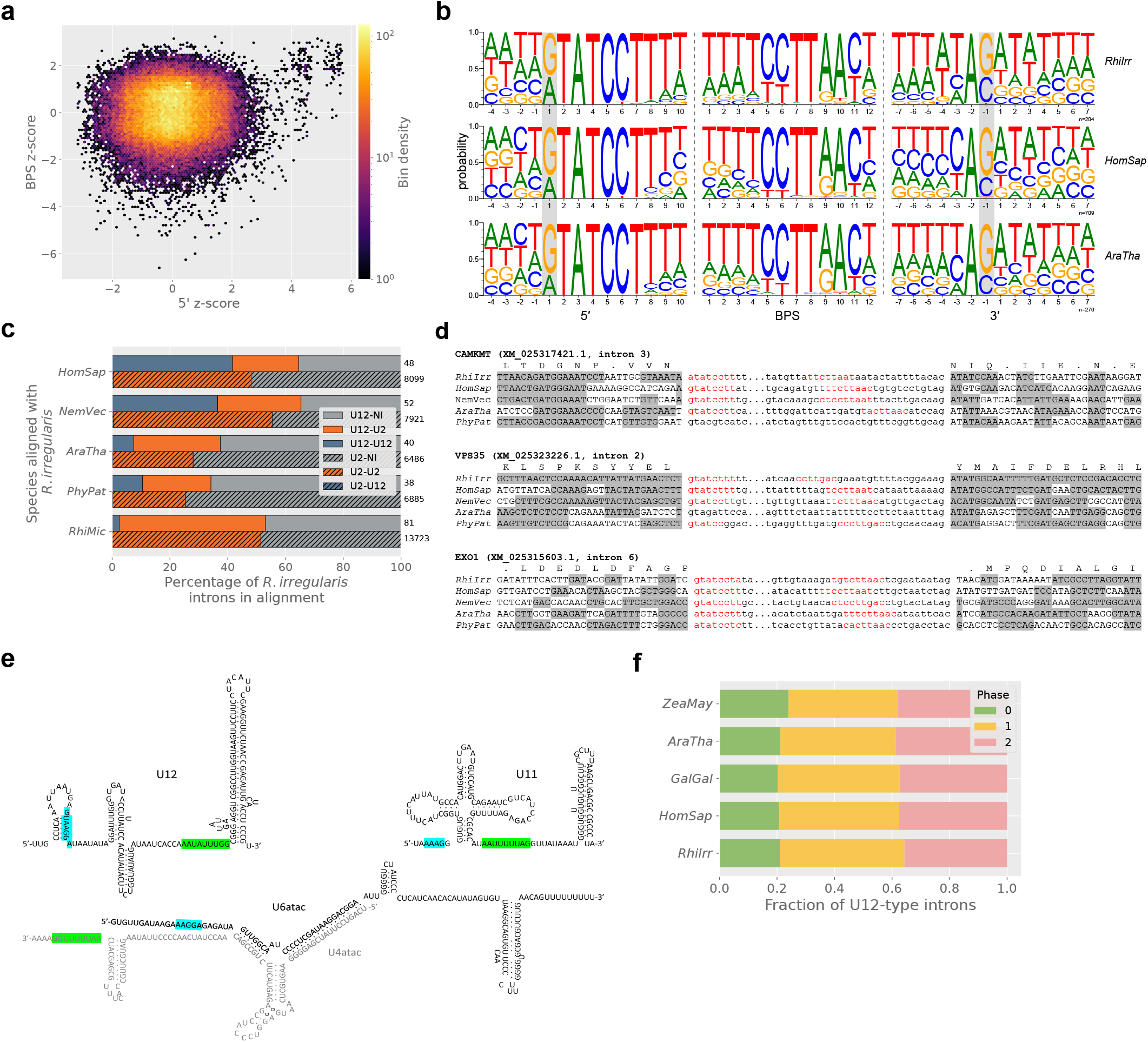
Evidence of minor introns and splicing machinery in *Physarum polycephalum*. (a) BPS vs. 5*′*SS scores for *Rhizophagus irregularis*, showing the expected cloud of introns with minor-intron-like 5*′*SS and BPS scores in the first quadrant. (b) Comparison of minor intron sequence motifs in *Rhizophagus*, human and *Arabidopsis*. (c) Conservation of *Rhizophagus* minor and major introns in different species. (d) Examples of minor introns in *Rhizophagus* in conserved alignments with minor introns in other species. (e) The four minor snRNAs U11, U12, U4atac and U6atac found in *Rhizophagus*. (f) Comparison of minor intron phase distributions in different species, showing the expected pattern in *Rhizophagus*. Species abbreviations are as follow: HomSap: *Homo sapiens*, NemVec: *Nematostella vectensis*, AraTha: *Arabidopsis thaliana*, PhyPat: *Physcomitrium patens*, RhiMic: *Rhizopus microsporus*, ZeaMay: *Zea mays*, GalGal: *Gallus gallus*.

### 3.11 No evidence for increased minor splicing in proliferating cells of R. irregularis

We next sought to test whether *R. irregularis*, like animals and plants, upregulates splicing of minor introns in proliferating cells. We used published transcriptomic data from five cell types (four replicates each), and assessed likely proliferation profiles of the six cell types using the previously published proliferation index (PI) approach. Briefly, we first identified putative orthologs of genes known to be associated with cell proliferation in humans. For each such putative PI ortholog, z-scores were calculated for all 20 samples, and those z-scores were then used for comparison across cell types as well as for comparisons within cell types between putative PI orthologs and other genes. This allowed us to calculate relative proliferation scores for all five cell types. While 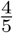 cell types showed similar PI values, one cell type, immature spores, showed substantially and significantly higher values (Fig. 12a), a pattern that also held when we look at the more straightforward metric of adjusted FPKM values (Fig. 13). This overall significance notwithstanding, it should be noted that only a small fraction of genes included in the PI individually showed significant differences in expression between cell types. In addition, we noted that many non-PI genes are also overexpressed in immature spores relative to other cell types; while one interpretation of this result is that it reflects generally more active gene expression in proliferating cells, it does provide a caveat for the overall strength of the observed difference.

**Fig. 12.**
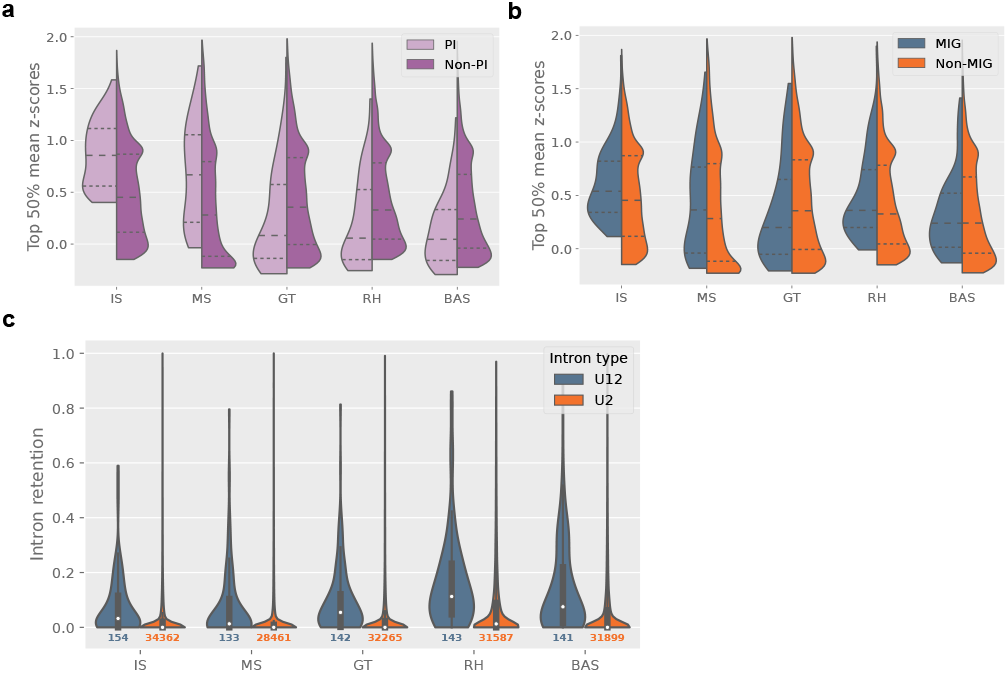
(a) Comparison of expression of proliferation-index genes (PI, light purple) and all other genes (Non-PI, dark purple) across cell types, n=70 PI and n=9276 non-PI in each cell type. (b) As in (a), but for minor intron-containing genes (MIGs) compared to non-MIGs; n=96 MIG and n=9249 non-MIG for each cell type. (c) Intron retention values across cell types for U12- (blue, left) and U2-type (orange, right) introns. Cell types are labeled as described in the text.

We then tested the association between markers of minor spliceosomal activity and these proliferation scores. We first looked for systematic differences in overall gene expression of MIGs between cell types with different proliferation scores, using various approaches. First, using the same z-score based approach as for the proliferation score (though with MIGs instead of putative PI orthologs), we found that MIGs were in fact more highly expressed in cell types with higher proliferation scores (Fig. 12b). On the other hand, we found that very few MIGs reached significant levels of differential expression, and were in fact underrepresented among genes that showed significant differential expression in multiple comparisons between cell types of different proliferation index scores (e.g., 4.2% of minor intron-containing genes compared to 21.9% of other genes in the IS-MS comparison). In total, these results suggest that expression of MIGs shows a detectable but only moderate association with proliferation index in *R. irregularis*, in contrast to the robust results previously observed in humans.

We next compared the efficiency of minor intron splicing between cell types. Contrary to our hypothesis that minor splicing would be more active in proliferating cells, we found that minor intron retention was in fact significantly (though only modestly) higher in proliferating cells (Fig. 12c). This result held whether we used z-score-based metrics or the intron retention values themselves, and whether we used splicing efficiency or intron retention as our metric. We also assessed expression of the minor splicing machinery itself (i.e., the known components of the minor spliceosome). In comparisons between immature spores and other cell types, no component individually showed higher expression, however collectively the machinery was 3.5x more highly expressed in immature spores than other cell types, reaching significance when considered collectively. However, the major spliceosomal machinery also showed a similar pattern (with 5x higher expression), and as such it seems that lower expression of the minor splicing machinery could be part of a larger pattern of up/regulation of core molecular functions in proliferating/quiescent cells.

The observed association between minor intron splicing and cell proliferation in animals resonates with the longstanding finding that minor introns are overrepresented in genes involved in core cellular processes. Given that minor intron splicing in *Rhizophagus* does not appear to be associated with cell proliferation, we probed these patterns more deeply.

Gene ontology analysis of *Rhizophagus* MIGs revealed a curious pattern in which GO results were highly dependent on the control dataset used. Because of the dearth of *Rhizophagus* functional annotations, GO analyses were necessarily run by identifying human orthologs of *Rhizophagus* MIGs. When GO analysis was run on these orthologs as a subset of all human genes, a number of overrepresented functional categories were found, in large part mirroring results for humans. However, we realized that there is a potential bias in this analysis: all human genes present in the *Rhizophagus* MIG ortholog set have *Rhizophagus* orthologs, thus excluding most human genes (only 14%, 3190/23257, had identified *Rhizophagus* orthologs), and in particular animal-specific genes. Remarkably, when we limited our GO analysis control group to human genes with *Rhizophagus* orthologs, we found much less functional overrepresentation (Table 4).

**Table 4.**
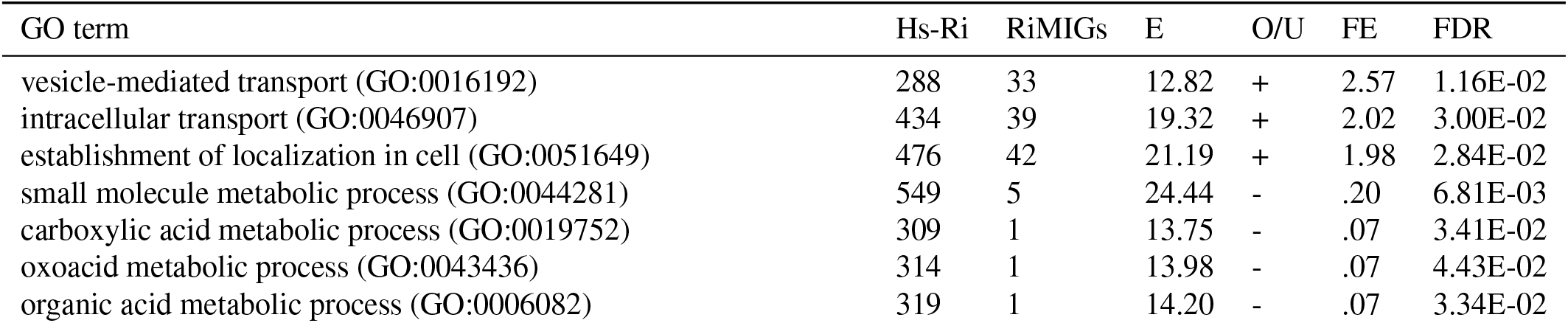
GO term enrichment for MIGs in *Rhizophagus* (RiMIGs), compared to all human-*Rhizophagus* orthologs (Hs-Ri). E: expected, O/U: over/under, FE: fold enrichment, FDR: false-discovery rate.

| GO term | Hs-Ri | RiMIGs | E | O/U | FE | FDR |
| --- | --- | --- | --- | --- | --- | --- |
| vesicle-mediated transport (GO:0016192) | 288 | 33 | 12.82 | + | 2.57 | 1.16E-02 |
| intracellular transport (GO:0046907) | 434 | 39 | 19.32 | + | 2.02 | 3.00E-02 |
| establishment of localization in cell (GO:0051649) | 476 | 42 | 21.19 | + | 1.98 | 2.84E-02 |
| small molecule metabolic process (GO:0044281) | 549 | 5 | 24.44 | - | .20 | 6.81E-03 |
| carboxylic acid metabolic process (GO:0019752) | 309 | 1 | 13.75 | - | .07 | 3.41E-02 |
| oxoacid metabolic process (GO:0043436) | 314 | 1 | 13.98 | - | .07 | 4.43E-02 |
| organic acid metabolic process (GO:0006082) | 319 | 1 | 14.20 | - | .07 | 3.34E-02 |

Notably, a similar concern applies to human MIGs in general: because nearly all human minor introns are quite old, human MIGs are commensurately old, which could drive functional correlations given known differences in functional categories between genes of different ages. Indeed, when we performed a GO analysis of human MIGs with *Rhizophagus* orthologs, limiting the reference set to human genes with *Rhizophagus* orthologs (a rough surrogate for gene age given that, unlike baker’s yeast, *Rhizophagus* may not have lost many ancestral genes [75]), we found a much lower degree of functional enrichment (Table 5). These results support the conclusion that the long-standing result that minor introns are functionally overrepresented in core cellular processes may be largely explained by the fact that minor introns fall primarily in evolutionarily older genes, which are overrepresented in core cellular functions. Interestingly, when we compared all human MIGs (given that minor intron presence strongly suggests that a gene is ancient) to human genes with *Rhizophagus* orthologs, we did see a significant number of overrepresented functional categories (https://doi.org/10.6084/m9.figshare.20483841). It is not entirely clear why all MIGs, but not MIGs with *Rhizophagus* orthologs, show substantial functional differences relative to all genes with *Rhizophagus* orthologs. Insofar as MIGs are ancient genes, MIGs without *Rhizophagus* orthologs likely represent losses in fungi; gene losses are likely to be functionally biased, perhaps explaining the observed pattern.

**Table 5.**
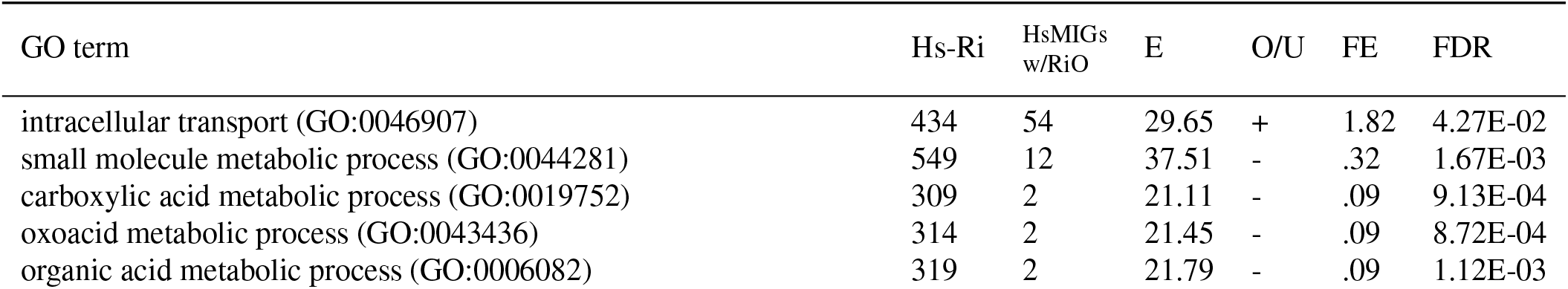
GO term enrichment for human MIGs with *Rhizophagus* orthologs (HsMIGS w/RiO), compared to all human-*Rhizophagus* orthologs (Hs-Ri). E: expected, O/U: over/under, FE: fold enrichment, FDR: false-discovery rate.

## 4 Discussion

### 4.1 An expanded view of minor intron diversity

Over the last decade (and after many if not all of the most prominent papers examining minor intron diversity were published), there has been a marked increase in the number of annotated genomes publicly available for bioinformatic analysis. Ten years ago, for example, NCBI had annotated fewer than 60 genomes—it now lists over 800, and that is counting only annotations performed by NCBI itself. The breadth of data now available has enabled us to undertake a much more sweeping, if necessarily less focused, assessment of minor intron diversity than has ever been possible before, uncovering a wide variety of both novel and confirmatory information about minor intron dynamics across the eukaryotic tree.

We have shown for the first time the presence of substantial numbers of minor introns as well as minor spliceosomal snRNAs in a variety of lineages previously thought to be lacking them, including green algae, fungi and stramenopiles. In addition, we have described findings contradicting a number of longstanding results in the minor intron literature, and have highlighted various underappreciated differences between MIGs and other genes while broadening the scope of earlier, more limited analyses.

Although we have endeavored to be as careful as possible in curating the data we have reported, as with most computational studies of this scale there is bound to be some noise, especially given our reliance on existing gene annotations derived from heterogeneous pipelines. One persistent issue in bioinformatic analyses of minor introns is the lack of a gold standard, empirically-verified set of minor intron sequences. While comparative genomics can do a great deal of heavy lifting in this regard, it is often a time-consuming process at scale and the field in general would benefit greatly from a ground-truth set of minor introns. We look forward to this type of data—based upon minor spliceosome profiling or another similar empirical method—being used to improve the accuracy of minor intron identification and as a result, furthering our understanding of minor introns and their evolutionary dynamics.

### 4.2 A complex history of minor intron evolution

These results underscore a complex history of minor intron evolution. We greatly expand the number of major eukaryotic groups known to contain minor introns including multiple unicellular lineages, highlighting the punctate distribution of minor introns. We show that multiple distantly-related lineages of fungi contain minor intron densities comparable to animals and plants. However, these three groups show dramatically different patterns of minor intron distribution. At one extreme, land plants show a high degree of minor intron stasis, with similar minor intron densities across nearly all studied plants. At the other extreme, fungi exhibit a wide diversity, with high minor intron densities in multiple lineages, greatly reduced numbers in multiple others, and complete absence from the globally dominant group Dikarya. Animals are somewhat intermediate, with minor intron presence in nearly all groups, but a range from very high to very low densities, and even multiple independent complete losses of minor introns. Of particular interest is the case of Dipterans, which exhibit massive reduction across the group, and yet almost no cases of complete loss. If is of great interest why these few minor introns have been so strongly retained across this clade. The diversity of minor intron conservation is also observed in terms of rates, with remarkable stasis in some groups (particularly vertebrates) contrasting with rapid turnover within single genera (e.g., *Blastocystis*, 92% intracellular transport (GO:0046907) 434 54 29.65 + 1.82 4.27E-02 small molecule metabolic process (GO:0044281) 549 12 37.51 - .32 1.67E-03 carboxylic acid metabolic process (GO:0019752) 309 2 21.11 - .09 9.13E-04 oxoacid metabolic process (GO:0043436) 314 2 21.45 - .09 8.72E-04 organic acid metabolic process (GO:0006082) 319 2 21.79 - .09 1.12E-03 (11/12) of minor introns in alignments between *Blastocystis sp. subtype 1* and *Blastocystis hominis* lost in *B. hominis*). We also document the remarkable diversity of the mechanisms by which minor introns are lost from genomes, ranging from nearly complete intron deletion to nearly complete conversion into major introns.

Our ancestral reconstructions suggest that ancestors of major groups (plants, animals, fungi) likely had modern densities comparable to the most minor intron-rich modern organisms (aside from the exceptional case of *Physarum* [41]). Coupled with very little evidence for *de novo* minor intron creation, this suggests a portrait in which modern organisms are largely minor intron-rich insofar as they have retained ancestral minor intron complements. The implied portrait contrasts with notions of multicellular organism has “highly-evolved”; rather, higher minor intron complements largely reflect lack of evolutionary change. The same contrast applies to within-kingdom comparisons: in particular, the animal lineages that have lost their minor spliceosomal systems (nematodes, myxozoa, oikopleuridae, tardigrades) have all been found to be generally fast-evolving at the genome level, including in terms of ancestral loss of spliceosomal introns overall ([46, 66, 55], GEL and SWR unpublished data).

### 4.3 Many features of minor intron evolution are consistent with neutral evolution

Attempts to make sense of the minor introns have generally alternatively argued they are functionally important or deleterious. Arguments for minor introns’ importance have noted their over-representation in genes with certain functions, associations of minor splicing with cell differentiation including apparent master regulatory roles, and one influential study finding greater evolutionary conservation of minor introns. Arguments that minor introns are deleterious tend to invoke their generally lower efficiency of splicing in addition to the complications and costs associated with maintaining two separate spliceosomal machineries.

Our results are not supportive of either of these perspectives as a general explanation for minor introns across eukaryotes. First, we report that apparent functional biases among minor intron-containing genes may be largely explained by minor introns bias towards ancient genes: because most minor introns are old, most MIGs are old, and core cellular processes are over-represented among old genes. This suggests that minor intron distributions across genes could simply reflect largely unbiased minor intron gain in ancestral genomes, followed by a lack of minor gain in more recent evolutionary time. Second, from our preliminary data in the minor intron-rich fungus *Rhizophagus irregularis* it does not appear that minor splicing is associated with cell proliferation in this species, suggesting that such an association may be specific to certain lineages and thus not capable of explaining general features of minor introns across eukaryotes. Third, we find a remarkably constant (though slight) trend for lesser, not greater, conservation of minor introns compared to major introns. Interestingly, we find very similar rates of intron loss by genomic deletion for minor and major introns, suggesting that minor introns’ somewhat lower overall evolutionary conservation reflects minor introns’ “extra” mechanism of loss through conversion to major introns. Our finding of similar rates of minor and major intron deletion is also not as predicted if minor introns are deleterious relative to major introns. Notably, this lack of an excess of minor intron loss is also observed in the lineages experiencing high degrees of minor-to-major conversion, which represent the best candidates for lineages in which minor introns might be deleterious. In total, then, our results suggest that neutral processes can explain much of the observed minor intron pattern across eukaryotes. This is not to say that all minor intron evolution is neutral, particularly in light of important cases of regulated and alternative splicing of minor introns; however, it may be the case that neutral processes govern most minor introns under most circumstances, and thus dominate the patterns across both genomes and lineages.

### 4.4 Secondary recruitment of ancient machineries for cell cycle regulation

Prior to the current work, we perceived a chicken and egg problem of functional biases among MIGs [11, 44] and control of cell proliferation by regulation of minor splicing ([27, 52, 2]: that is, how could the regulatory control evolve without the functional bias, by why would the functional bias evolve without the regulatory function? We thus sought to illuminate this question by studying a third minor intron-rich lineage. The current findings that the observed functional biases appear to be largely explained by minor introns’ bias towards older genes, and older genes bias towards core cellular functions, suggest an answer. Thus, functional biases could have initially evolved due to these gene age biases, and this functional bias could then have secondarily been recruited to regulate cell proliferation in animals in plants.

While this scenario makes sense schematically, is remains a remarkable contention that decreased minor splicing could evolve a function in cell regulation; insofar as MIGs represent a quasi-random subset of ancient genes, it seems likely that a global reduction in minor splicing would have a wide variety of impacts, many of them likely costly. Thus how failure to process a quasi-randomly chosen set of ancestral genes could evolve as a regulatory mechanism remains puzzling, and will require additional work across diverse minor intron-containing lineages.

Our results do not support the emerging dominant hypothesis for the existence of minor introns, namely that minor introns provide a means for regulation of cell cycle progression. The reported lack of cell cycle-regulated minor intron splicing in fungi suggests that this association is not a general phenomenon, correspondingly weakening the hypothesis that such a function could explain the persistence of minor introns across eukaryotes generally. However, given the possibility that it may be fungi that are atypical, having secondarily lost this function, discovery and study of additional minor intron-rich lineages is a priority, as is development and testing of alternative hypotheses for the origins and functional biases of minor intron-containing genes.

### 4.5 Limitations of the *Rhizophagus* analysis

Possible caveats of this analysis arise from two surrogates that we have employed. First, to assess cell cycle activity/proliferation of cell types, we have used orthologs of human genes associated with proliferation. The possibility of turnover of gene expression patterns raises the concern that these genes are not an appropriate gene set to assess proliferation. Indeed, while clear statistical differences in proliferation are seen when PI genes are viewed collectively, only a small fraction (∼5-10%) individually show significantly different expression between cell types. However, similar comparisons with model fungi attest to generally good conservation of genes’ association with proliferation, consistent with an ancient core of cell cycle regulation. Second, we have used available transcriptomic data not specifically generated for the purposes of comparing proliferation, potentially leading to noise in the data. However, the general pattern observed, in which developing spores show the highest proliferation index, mirrors intuitive expectations, suggesting that our proliferation scores are capturing at least some of the relevant biological phenomena. Testing of transcriptomic effects of direct manipulations of cell cycle would be very useful to confirm (or refute) our results.

## 5 Data availability

Plain text data for Fig. 1 is archived on FigShare at https://doi.org/10.6084/m9.figshare.20483655. Additional versions of certain figures, including a rectangular version of Fig. 1 and interactive versions of the plots in Fig. 9 are available at https://github.com/glarue/minor_introns.

## 7 Supplementary materials

### 7.1 Supplementary figures

**Fig. 13.**
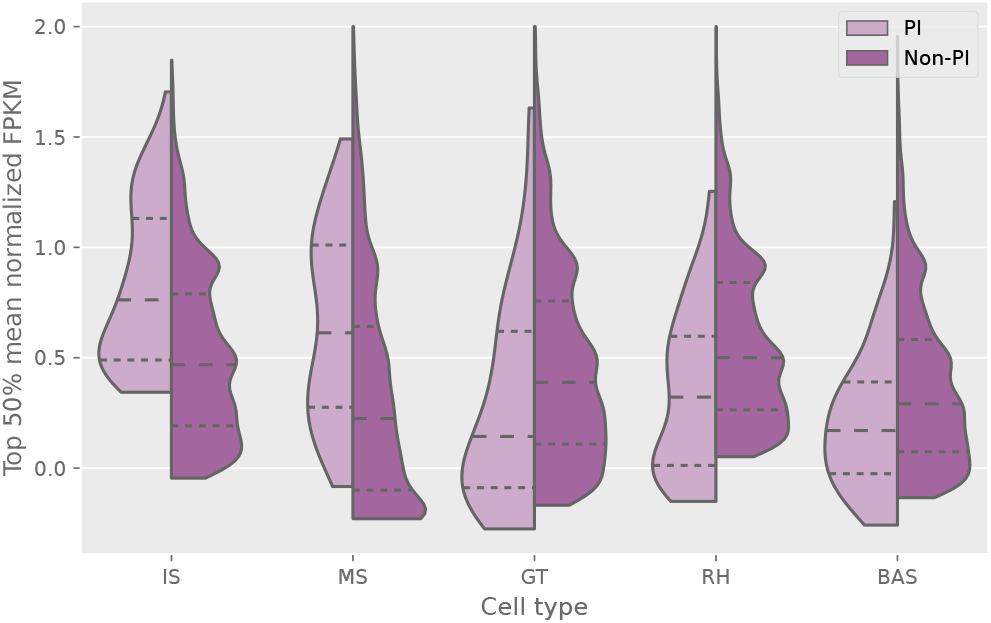
Comparison of expression (normalized FPKM from `DESeq2`, power-transformed) of proliferation-index genes (PI, light purple) and all other genes (Non-PI, dark purple) across cell types.

## Footnotes

1 Incorrectly labeled in the NCBI database as *Planoprotostelium fungivorum*; originally described as *Planoprotostelium fungivorum* in [31] but subsequently corrected in the main text (the incorrect usage remains in the supplemental materials); see [78] for supporting evidence of its classification as *Pr. aurantium*. Credit to Marek Eliáš for this addendum.

2 We note here for posterity the most striking case of this bias we have found in the yeast species *Candida maltosa* (which lacks minor introns), where all ∼1000 annotated major introns appear to be phase 0 (Fig. 7c, bottom-right corner).

